# Kinetic Proofreading through Parallel Reactions on a Single T Cell Receptor

**DOI:** 10.1101/2025.06.01.657268

**Authors:** Shumpei Morita, Jay T. Groves

## Abstract

T cells can recognize a few molecules of cognate antigen amongst vastly outnumbering non-cognate ligands. The T cell receptor (TCR) differentiates antigens based on antigen-TCR binding dwell time through a kinetic proofreading process. Historically, this has been modeled as the ligated receptor undergoing a series of reactions before producing a signal. In such a sequential mechanism, the number of steps is a key determinant of discrimination fidelity. Here, we consider two features of the molecular mechanism of TCR activation that diverge from a sequential process and suggest that an alternative kinetic proofreading mechanism may be at play. First, activation processes of multiple ITAM domains of the TCR represent parallel reaction sequences taking place on a single TCR molecule. Second, the states of the parallel proofreading reactions are integrated to produce a binary output from each TCR in the form of a discrete LAT condensation event, which may or may not occur. We examine a revised kinetic proofreading scheme based on parallel reactions followed by an integration step (multi-thread scheme) and compare its performance with the sequential scheme in a stochastic setting. A distinct difference in a multi-thread scheme is that multiplicity of the parallel reaction threads provides an additional means to increase discrimination fidelity. This relieves the need for fine-tuned kinetics among chemically distinct reaction steps, which is a major hurdle for physical implementation of a sequential mechanism. Lastly, we reinterpret previously reported experimental observations and find that various proofreading behaviors are well described as proofreading through parallel reaction threads.

**Significance Statement:** The kinetic proofreading mechanism by which T cells discriminate antigen has attracted interest from both physical and immunological perspectives. The proofreading process has generally been modeled as a multi-step sequence of reactions, and many experimental observations have been interpreted based on this presumed mechanism. However, these models do not capture key molecular features of the TCR signaling mechanism, including ITAM multiplicity and LAT condensation. Here, we find distinct consequences of these molecular features by examining kinetic models and reinterpreting published experimental data. ITAM multiplicity offers multiple parallel reaction threads, re-integrated by the LAT condensation step, which readily improves discrimination fidelity up to the observed levels. Multi-thread reactions are suggested to play a central role in amplifying TCR kinetic proofreading performance.

## Introduction

T cells detect rare agonist peptide-major histocompatibility complex (pMHC) molecules dispersed among abundant self pMHC on antigen presenting cell (APC) surfaces. This remarkable antigen discrimination capability is achieved through a kinetic proofreading mechanism, in which T cell receptor (TCR) activation depends on the binding dwell time of pMHC molecules (1–3). The mean dwell time difference between agonist-and self-pMHC is only about 10-fold, while the abundance difference can exceed 1000-fold (2–5). T cells also exhibit extreme sensitivity; a few tens of agonist pMHC:TCR binding events are sufficient for T cell activation (6–10). The small number of binding events ensures high levels of stochastic variation, which puts additional limits on the ligand sensing accuracy (11–15). Kinetic proofreading by TCR, when properly conditioned, is thought to provide sufficiently high discrimination fidelity to successfully recognize antigen in this challenging context (16).

Although TCR kinetic proofreading is a well-established concept, how the molecular signaling mechanism achieves kinetic proofreading remains unclear. The TCR proofreading mechanism has mostly been modeled as a Markov process based on sequential reactions (here referred to as the sequential scheme, **Fig. 1A**). Upon ligation, the receptor progresses through a series of intermediate steps before a signaling active state is reached. If the ligand unbinds before the receptor reaches its activated state, it quickly reverts back to the fully deactivated starting state and no signal is produced. Intermediate reactions in the TCR activation process include phosphorylation of ITAM domains on the TCR by the kinase Lck, recruitment of the Zap70 kinase to phosphorylated ITAMs, and activation of Zap70 through phosphorylation by Lck (17, 18). Once activated at the TCR, Zap70 begins to phosphorylate LAT molecules. Phosphorylated LAT get crosslinked by Grb2, SOS, and other molecules, ultimately forming a type of signaling protein condensate, which then activates downstream pathways (18–24). Lck recruitment, ITAM phosphorylation, Zap70 activation, and TCR-proximal LAT phosphorylation have all been experimentally implicated as possible proofreading steps (25–30). However, inconsistencies exist among reports and the identities of the most critical proofreading steps remain uncertain. Only sufficiently slow steps can effectively participate in kinetic proofreading. Thus, without detailed information on *in situ* reaction kinetics it is essentially impossible to determine from the reaction sequence alone which steps are effective contributors to proofreading. Nonetheless, several reports have examined experimental measurements of T cell kinetic proofreading fidelity to infer the effective number of proofreading steps based on the sequential scheme (16, 27, 31, 32). Other works have considered sequential reaction schemes with modifications such as extended feedback/feedforward pathways to describe further details of T cell response characteristics as well as mutation effects (30, 33–36).

**Figure 1.**
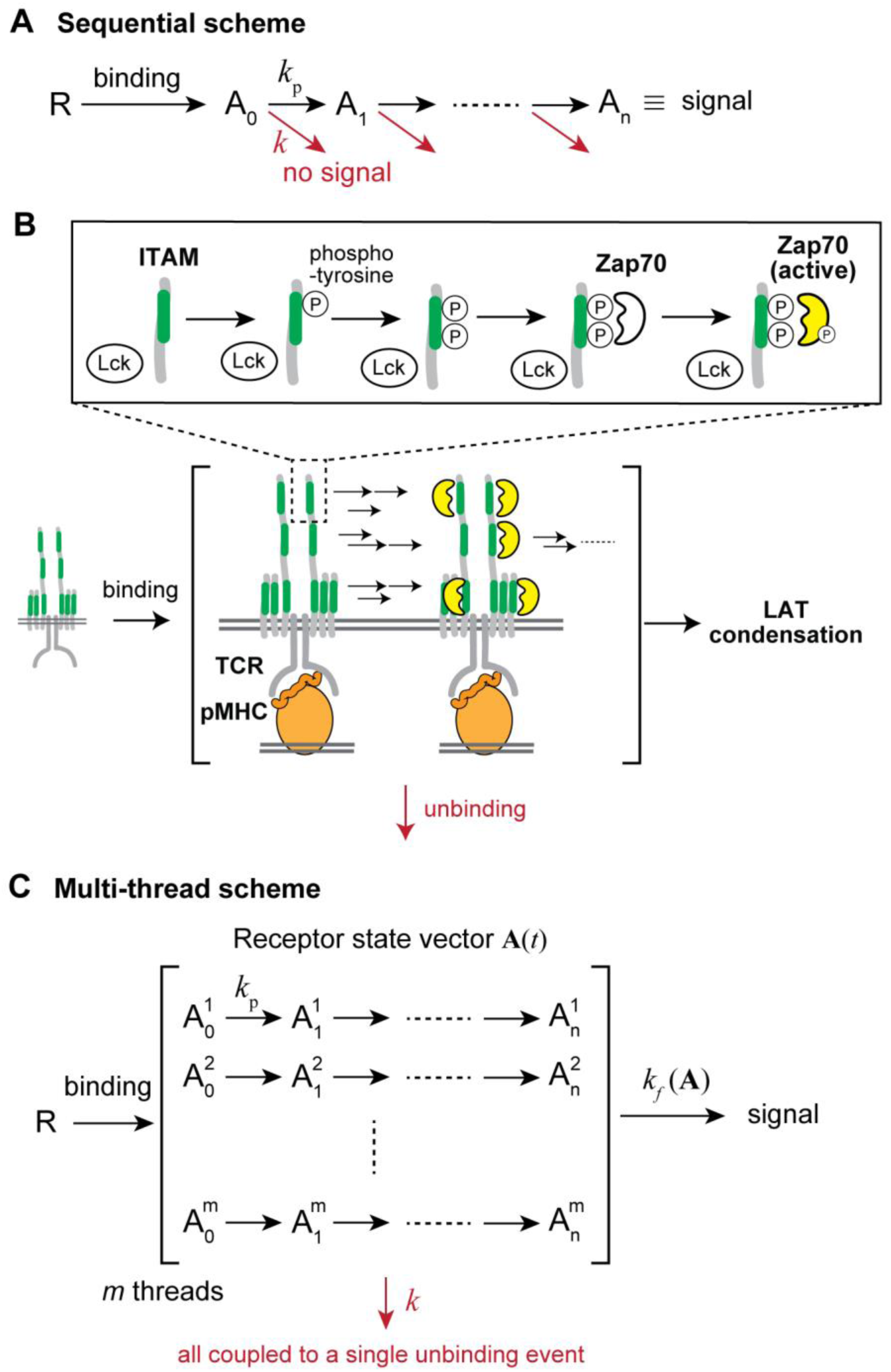
Construction of revised kinetic proofreading scheme reflecting ITAM multiplicity and binary LAT condensation. (**A**) Diagram of sequential scheme. (**B**) Schematic representation of the parallel signaling at multiple ITAM domains followed by LAT condensation. (**C**) Diagram of multi-thread scheme.

In the present work, we examine how two features of the molecular mechanism of TCR activation that do not follow a sequential activation process can affect kinetic proofreading. The first feature is ITAM multiplicity. There are ten ITAM domains within TCR:CD3 cytoplasmic tails, each of which can recruit and activate a Zap70 molecule during TCR ligation (17, 37). Upon ligand binding, multiple proofreading reaction sequences are initiated on the individual ITAM domains of the same receptor. The second feature relates to how an activated TCR drives LAT condensation. Recent observations have revealed that a single pMHC:TCR complex can nucleate a discrete LAT condensate in its proximity (10, 38), and each LAT condensate that successfully forms produces a cell-wide calcium spike within seconds (39). These LAT condensation events represent binary quanta of signaling output from individual TCR and provide an integration mechanism that recombines outputs from the parallel reactions on multiple ITAM domains of the receptor (**Fig. 1B**). We incorporate these features in a model of kinetic proofreading as a “multi-thread scheme”, in which multiple sequences of proofreading steps (here referred to as threads) occur in parallel on a single ligated receptor (**Fig. 1C**). The ensemble of reaction threads, corresponding to the states of individual ITAMs and associated Zap70 molecules, are simultaneously initiated by binding of ligand to the TCR. Each ITAM reaction thread then progresses independently through sequential activation steps leading to Zap70 activation. The resultant Zap70 kinase activity from multiple ITAM threads is re-integrated by the LAT condensation step, which provides a binary, all-or-nothing, output at the individual receptor level.

We computationally compare performance characteristics of sequential and multi-thread proofreading schemes in a stochastic setting. The results reveal that, for the sequential scheme to achieve sufficient discrimination fidelity, many proofreading steps are necessary. Moreover, the rates of individual reactions must be similar in order to effectively contribute towards proofreading fidelity. In general, only a subset of rate-limiting steps will dominate the behavior and degrade the performance. For example, 10-step reactions with 20-fold rate heterogeneity provide only about 3 effective steps in a sequential mechanism. By contrast, the multi-thread scheme can achieve enhanced discrimination fidelity by having more threads, nearly multiplying the effective number of steps by the number of threads. Since the individual threads are essentially chemically identical copies of the basic reaction sequence, their rates are intrinsically uniform. We postulate that such a multi-thread scheme is more evolutionarily accessible, requiring only domain multiplicity without need for finely matched kinetics from distinct chemical reactions. Lastly, we re-analyze a variety of experimental data in terms of a multi-thread proofreading scheme, revealing that the TCR behaviors which have been described as proofreading through sequential reactions are readily re-interpreted as proofreading through parallel threads. Mechanistically, the results suggest that as few as one rate-limiting step in the reaction sequence occurring on several ITAMs is sufficient to describe experimentally measured antigen discrimination fidelity. ITAM multiplicity, and the resultant multi-threaded reaction mechanism on TCR, thus appears to be the primary source of proofreading fidelity.

## Construction of a multi-thread kinetic proofreading scheme

### Kinetic schemes for individual ligand binding events

We construct the sequential kinetic proofreading reaction scheme following earlier studies such as (1). Upon ligand binding, the receptor undergoes multiple reaction steps (*n* steps) with a forward reaction rate *k*_p_ until the final state is reached (**Fig. 1A**). Ligand unbinding can occur throughout the process with an off-rate *k*. If the final state is reached before ligand unbinding, a binary signal is produced, and if ligand unbinding occurs before the final state is reached, no signal is produced.

For the multi-thread scheme, upon ligand binding to a single receptor, multiple sequences of multi-step reactions (threads) are synchronously initiated. Each thread then progresses stochastically and independently during the ligand dwell time (**Fig. 1C**). These threads represent parallel reactions at multiple ITAM domains of a single TCR. Threads are re-integrated at the final signal-producing step (here referred to as the firing step), which is physically associated with LAT condensation. The firing step occurs stochastically during the ligand dwell time, and its rate is determined by the number of activated Zap70 molecules, corresponding to the number of completed reaction threads, on the TCR. The firing rate, which corresponds to the experimentally measured propensity function for TCR signal activation (38), is time dependent. Ligand unbinding (off-rate *k*) can occur at any point in the process, which terminates the entire process. If firing occurs before unbinding, a signal is produced, and no signal is produced otherwise. Formally, we model the TCR to have *m* reaction threads, each of which consists of *n* sequential steps with rate constants *k_p_*. Each thread progresses through *n* + 1 states with a transition rate *k_p_*, and its time evolution follows a simple Markov process in the same manner as the sequential scheme. The state of a ligated receptor is represented by the ensemble of independent thread states, ***A**(t)*. The firing rate is expressed as a function of this state vector as *k_f_(**A**)*.

### LAT condensation non-linearly integrates multiple thread states into a singular firing event

LAT condensation nucleation occurs as a type of phase transition once some minimal number of nearby and multiply phosphorylated LAT molecules become mutually crosslinked by Grb2, SOS, and other proteins (22, 40). The ensuing LAT condensates grow rapidly (38) and mature to produce a cell-wide calcium spike within 10 s (39). LAT is phosphorylated (pLAT) by activated Zap70 on ligated TCR, and the momentary probability of LAT condensate nucleation (firing rate) is dependent on the number of activated Zap70 molecules on the TCR. Experimentally, LAT condensates are observed to persist even after ligand unbinding from TCR and neither the size nor the lifetime of the LAT condensate depends on the originating pMHC:TCR dwell time (38). LAT condensates represent discrete quanta of signaling output from individual TCR. Experiments further reveal a long, and broadly distributed, delay time between initial pMHC:TCR binding and LAT condensate nucleation. During this time, no measurable buildup of pLAT is observed, suggesting that high levels of phosphatase activity quickly dephosphorylate LAT. These observations indicate a plausible nucleation model in which pLAT levels in proximity to activated Zap70 on TCR fluctuate as a result of stochastic kinase-phosphatase competition (40–43). Once a pLAT density fluctuation exceeds a certain critical level, the nucleus stabilizes itself and grows abruptly (**Fig. 2A**). Candidate mechanisms for stabilization include phosphatase exclusion or positive-feedback regulation of kinases (21, 44). Different number of active Zap70 molecules on TCR, from zero to ten, is expected to change the fluctuation level and hence the momentary probability of nucleation.

**Figure 2.**
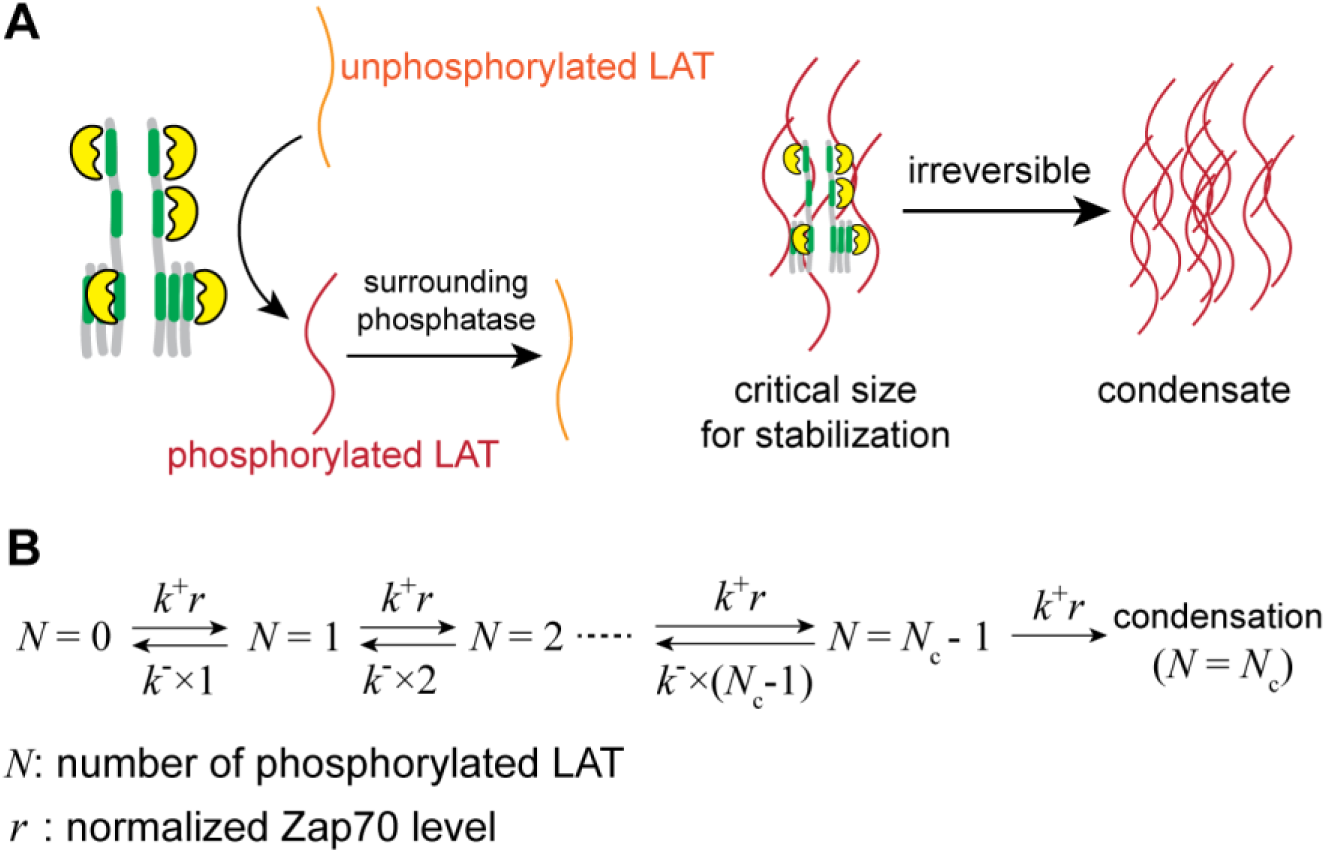
Modeling of LAT condensate nucleation kinetics. (**A**) Schematic of the fluctuation-driven LAT condensate nucleation model. (**B**) Reaction scheme representing the LAT phosphorylation fluctuations and condensation.

We construct the master equation for LAT condensate nucleation (firing) at TCR based on the above molecular mechanism (see **Fig. 2B** and **Appendix 1**). An analytical expression for the nucleation rate, *k_f_*, as a function of normalized Zap70 activity level, *r*, reveals power-law scaling with *k_f_* ∝ *r^Nc^*, where *N_c_* is the critical number of phosphorylated LAT tyrosine residues necessary for nucleation. LAT condensate nucleation by multiple Zap70 molecules is highly cooperative, specifically meaning that the nucleation propensity non-linearly scales with the local Zap70 level.

In the context of modeling the multi-thread scheme based on molecular details of nucleation, a Zap70 activation event is represented by the completion of each thread. Zap70 activity level *r* corresponds to the normalized count of completed threads, 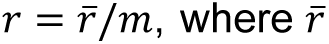 is the count of the threads in the final state. The nucleation rate corresponds to the firing rate, defined as

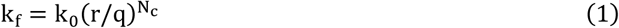

where *q* > 0 is an adjustable parameter representing the firing threshold. *k*_0_ is a constant, which will be used as an off-rate of agonist ligand (see the next subsection); it represents the time scale of the entire system in our model, which is why it is used here as a constant. We refer *N_c_* as the cooperativity exponent in the context of the multi-thread scheme. For simpler analyses, we also consider the large *N_c_* limit (referred to as sharp firing regime),

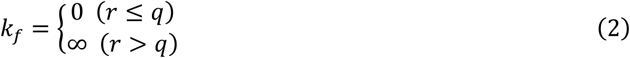

meaning that firing occurs immediately and deterministically once *r* exceeds *q*. In a sharp firing regime, *q* has redundancy between possible *r* values, and is chosen so that 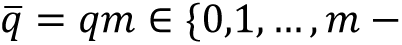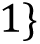 for simpler expression.

We note that LAT condensate nucleation is a highly complex process with active modeling efforts (40), and the model above is highly simplified to ignore factors such as molecular diffusion, LAT tyrosine multiplicity, or substrate depletion. Since these components may affect the resulting nucleation rate to some extent, we avoid extending our interpretations beyond a minimal qualitative conclusion that firing is cooperative.

### Cellular integration

We model the cellular signal integration from individual binding events based on our previously reported experimental observations (**Fig. 3A**). In single-molecule imaging studies that observe individual agonist pMHC binding and unbinding events, the bound pMHC:TCR complexes were well separated from each other (8, 9). An individual pMHC:TCR, when the dwell time is long enough, stochastically nucleates an individual discrete LAT condensate (38). LAT condensates additively induce calcium signal, which then induces NFAT translocation in a binary manner, marking early T cell activation (39). These events occur within several minutes, and NFAT translocation is observed when the cell has experienced about 50 binding events of agonist pMHC (9). These observations suggest a simple cellular integration mechanism, in which calcium spikes from a sequence of binding events (and corresponding LAT condensates) add, and a cell-wide response occurs if a threshold is crossed (**Fig. 3B**).

**Figure 3.**
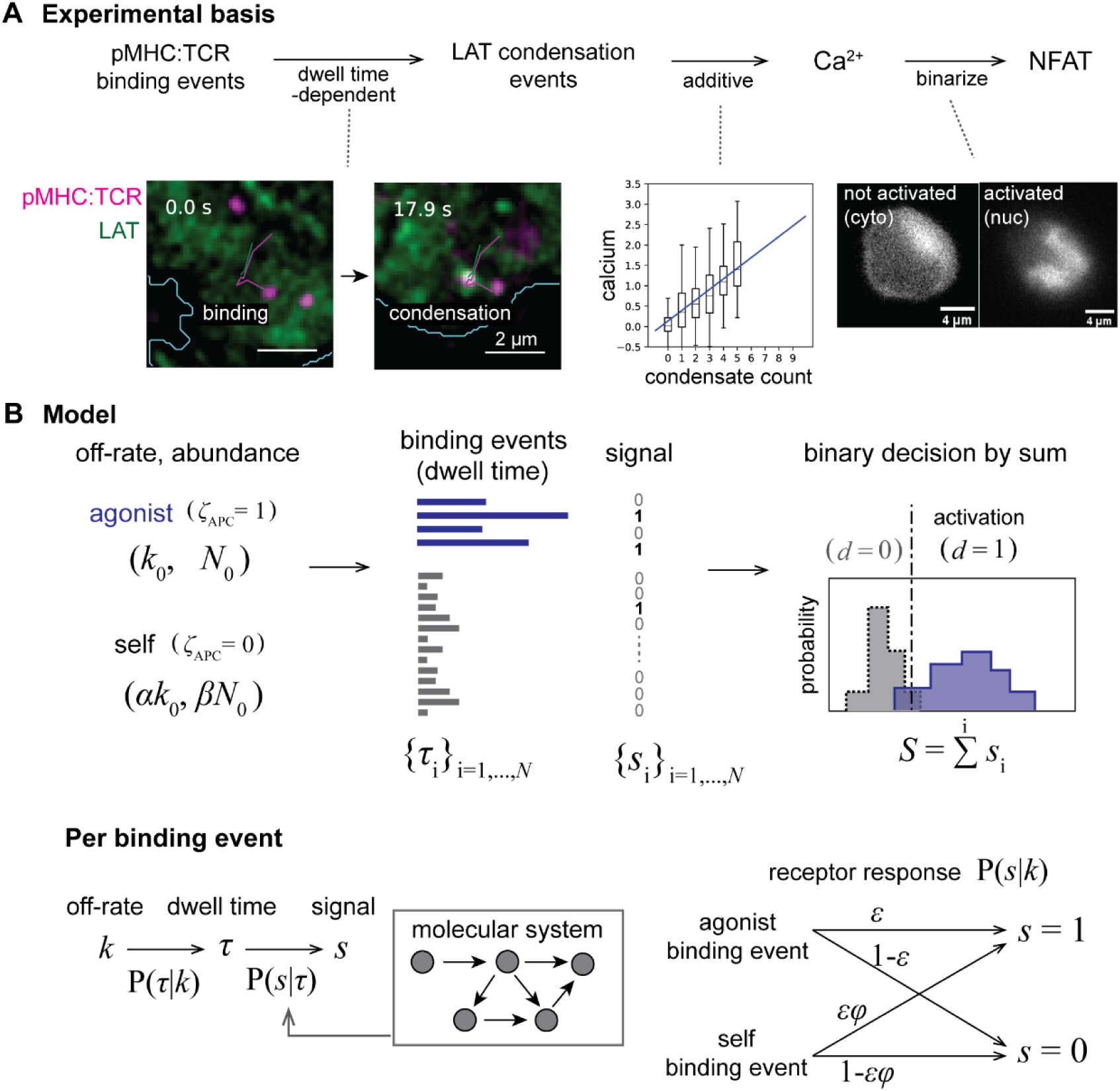
Model formulation based on experimental observations. (**A**) Schematic of experimentally observed T cell signaling cascades (38, 39). Images and plots are re-plotted from (39). (left bottom) Images of single-molecule pMHC:TCR binding events (magenta) and delayed proximal LAT condensation (green). (center bottom) Additive calcium response per discrete condensates. (right bottom) Images of binary activation of NFAT. (**B**) Cell-level model of antigen discrimination consistent with the experimental basis.

To formally define this, we suppose the task of binary classification, where APC can present either self pMHC (ζ_APC_ = 0) or agonist pMHC (ζ_APC_ = 1). Each type of ligand possesses two parameters, off-rate k and mean number of binding events ⟨*N*⟩. We set *k* = *k*_0_ for agonist and *k* = α*k*_0_ for self with the off-rate ratio α = 10. We also set ⟨*N*⟩_agonist_ = *N*_0_ and ⟨*N*⟩_self_ = β*N*_0_, where *N*_0_ = 50 and β = 1000. That is, a cell needs to respond to about 50 binding events of agonist, while ignoring about 50000 binding events of self with 10-times faster off-rate (see **Appendix 2** for more detailed justifications). Binding events are represented by the sequence of dwell times {τ_*i*_}_*i*=1,…,*N*_, where the actual number of binding events *N* follows a Poisson distribution with mean ⟨*N*⟩. Each dwell time τ follows an exponential distribution with off-rate *k* as P(τ|*k*) = *ke*^−*k*τ^. TCR-proximal signaling converts the dwell time sequence to binary signal sequence {*s*_*i*_}_*i*=1,…,*N*_, where no interference among binding events is considered, reflecting the confined pLAT proximal to individual pMHC:TCR due to phosphatase pressure. The cellular binary decision of activation *d* is made based on the summed signal *S* = ∑_*i*_ *s*_*i*_ as *d* = 1 for *S* > *S_c_* and *d* = 0 for *S* ≤ *S_c_*, where *S_c_* is a certain threshold value. In this model, the T cell acts as a binary classifier from ζ_APC_ to *d* with an adjustable paramter *S_c_* which determines false-positives and false-negatives, thus the separation of the probability distribution of *S* for each ligand type will determine the overall performance.

## Analysis of the models

### Strategy

The TCR-proximal molecular systems, such as those represented by sequential or multi-thread schemes, determine the probability of producing a binary signal given a certain dwell time P(*s*|τ). The probability of producing a signal per binding event given a certain off-rate can be determined as P(*s*|*k*) = ∫ *P(s*|τ)P(τ|*k*)*d*τ, which is here referred to as receptor response. As we consider two ligand types, receptor response is fully expressed by two values: receptor sensitivity ɛ = P(*s* = 1|*k* = *k*_0_) and error rate ϕ = P(s = 1| k = αk_0_)/P(*s* = 1|*k* = *k*_0_) (**Fig. 3B** bottom right). Since the number of binding events follows a Poisson distribution, the receptor response determines the probability distributions of cellular sum signal *S* as

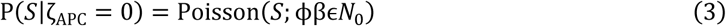

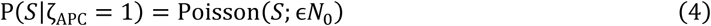

, where Poisson(⋅; *x*) denotes Poisson distribution with mean *x*. Here we can see that the receptor response (ϵ, ϕ) acts as the central parameter. For the molecular system of interest, we aim to deduce the (ɛ, ϕ) pair representing the receptor response. Once (ɛ, ϕ) is known, the antigen discrimination performance can be determined independently of specific molecular systems. We first analyze the latter by mapping the antigen discrimination performance in (ɛ, ϕ) space as preparation for analyzing specific molecular systems in the following.

### Antigen discrimination performance in a stochastic regime

We introduce the channel capacity as a metric of antigen discrimination performance. If agonist produces more sum signal *S* than self and the two distributions are separated well, binary classification can be performed well by having a proper threshold *S_c_*. Channel capacity can be used as a scalar metric of this separation (30), with the definition 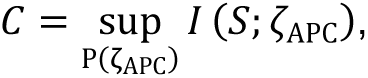 where P(ζ_APC_) is prior probability and *I(S*; ζ_APC_) is mutual information. Channel capacity represents the maximum information transmitted through the system with a range of 0 to 1 bit, and here we call it discrimination performance.

Discrimination performance *C* was mapped in (ϵ, ϕ) space (**Fig. 4A, B**). It decreases both for high error rate ϕ and low receptor sensitivity ɛ. The former is due to the loss of selective response. As ϕ increases up to β^−1^ = 10^−3^, *S* from self and agonist approach the same and become indistinguishable (compare top and middle panels of **Fig. 4A**). The latter is the result of stochastic noise. *S* becomes small for low ɛ and the stochastic noise (scale of *S*^1/2^) becomes prominent. For very small ɛ, agonist pMHC binding events produce zero signal (*S* = 0) with a significant probability and the performance is poor for any ϕ, reflecting inevitable large false-negatives (bottom panel of **Fig. 4A**). Analytically, compensation relationships can be seen (**Eq. 3, 4**): larger abundance ratio β requires lower error rate ϕ, and fewer binding events *N*_0_ requires higher receptor sensitivity ɛ. In the deterministic limit of *N*_0_ → ∞, ɛ no longer matters, and the perfect discrimination is achieved as long as ϕ < β^−1^. When stochasticity is considered, receptor sensitivity must be larger than about *N*^−1^, which puts an additional bound on the receptor response. As a result, the limited regime with low ϕ and high ɛ provides sufficient discrimination performance (**Fig. 4B**).

**Figure 4.**
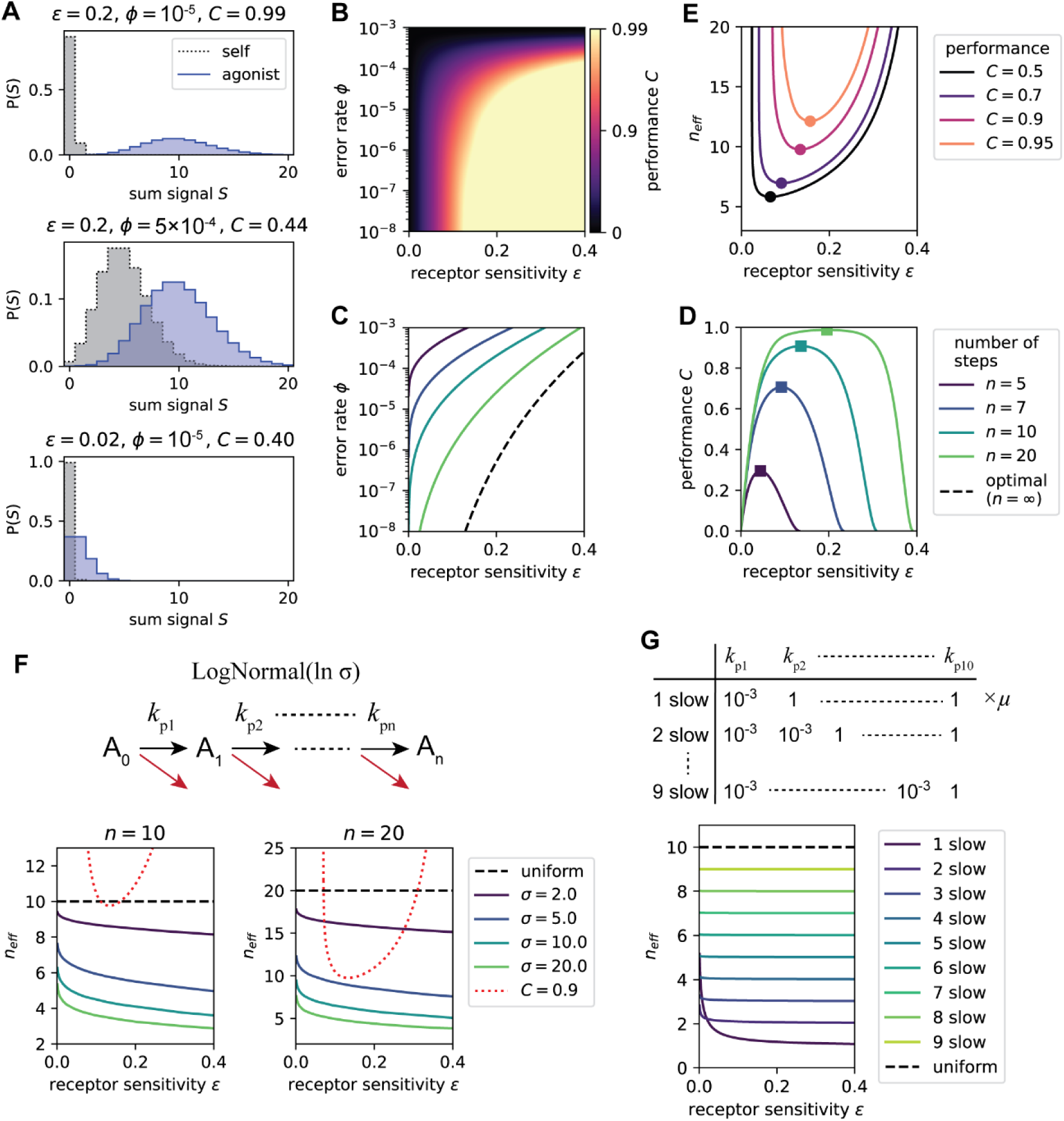
Performance of sequential scheme. (**A**) Representative sum signal distributions and resulting discrimination performance metrics. (**B**) Discrimination performance metric mapped in receptor response space. (**C**) Receptor response trajectories for different number of steps. (**D**) Discrimination performance metric corresponding to the trajectories in **C**. (**E**) Effective number of steps corresponding to the contours in **B**. Maximum and minimum are highlighted by markers in **D**, **E**. (**F**) The effect of heterogeneous forward reaction rates on the effective number of steps. Stochastic sampling from log-normal distributions with different relative variations were performed. Median values from multiple trials are shown. The curve for a benchmark *C* = 0.9 is overlaid. (**G**) The cases with defined number of slow steps.

In the following analysis, we primarily use *C* = 0.9 as a benchmark for good discrimination performance for clarity of description, but we will confirm that the conclusions do not depend on this arbitrary choice. This benchmark approximately keeps both false-positives and false-negatives below 5% with an appropriate choice of *S_c_* (**Fig. S1**).

### Sequential scheme requires many proofreading steps

Evaluation of molecular systems can be done by obtaining (ɛ, ϕ) pairs and looking up the corresponding discrimination performance. We first evaluate the sequential scheme with *n* proofreading steps and forward reaction rate *k_p_* (**Fig. 1A**). Treating *n* and *k_p_* as paramters, we get the trajectories of (ϵ, ϕ) achievable by this scheme. Since 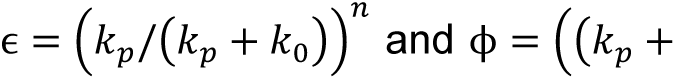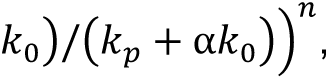 the trajectory follows

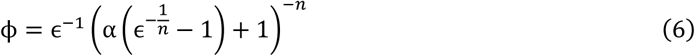

where the parameter *k_p_* moves the coordinates on this trajectory for a given *n*. The trajectories for several *n* are shown in **Fig. 4C**. Greater receptor sensitivity ϵ entails higher error rate ϕ, exhibiting a form of speed-accuracy trade-off (45–48). As a result, the discrimination performance metric exhibits bell-curves, where too low and too high ϵ (hence *k_p_*) exhibits poor discrimination performance (**Fig. 4E**). As shown in **Appendix 3**, ϕ monotonically decreases with *n*, covering the entire range from ϕ = 1 with *n* → 0 to the lowest possible ϕ with *n* → ∞. Thus, larger *n* always achieves better performance when ɛ (hence *k_p_*) is optimized. The monotonic improvement of performance over *n* makes the sequential scheme a useful ruler for comparison with other schemes discussed later. We define the effective number of steps *n*_eff_(ϵ, ϕ) as an inverse function: for a given (ϵ, ϕ), how many steps are necessary to achieve it by sequential scheme. The effective number of steps required for certain levels of discrimination performance are drawn in **Fig. 4D**. For the performance of *C* = 0.9, about 10 steps are required with optimal ϵ.

### Sequential scheme collapses to a subset of rate-limiting steps

We supposed uniform forward reaction rates in the analysis above. The actual reactions possess heterogeneity in rates, including both inherent factors owing to different biochemical reactions and stochastic factors such as protein expression levels. We evaluated heterogeneity effects by sampling forward reaction rates from log-normal distribution with mean rate μ and relative variation σ (see **Methods**). The obtained *n*_eff_ for several σ are shown in **Fig. 4F**. The effective number of steps was susceptible to the rate heterogeneity. To achieve *C* = 0.9, about 10 steps with nearly identical rates or 20 steps with high uniformity (σ < 5) are required. This performance degradation occurs because only a slow subset of steps will affect the term {*k_pi_*/(*k_pi_* + *k*)}. To confirm this understanding, cases with defined numbers of slow rate-limiting steps were analyzed (**Fig. 4G**). The effective number of steps indeed matched the number of rate-limiting steps.

### Limited performance of sequential scheme is unique to a stochastic regime

The requirement of many proofreading steps is unique to a stochastic regime. At *k_p_* → 0 limit (or ϵ → 0), error rate scales as ϕ → α^−*n*^. Perfect discrimination is achieved in deterministic regime when ϕ < β^−1^, which requires only *n* > 3. In a stochastic regime, both conditions ϕ < β^−1^ and ɛ > *N*^−1^ need to be satisfied, which increases the required number of steps. The susceptibility to rate heterogeneity can also be escaped in deterministic limit, because the same scaling ϕ → α^−*n*^ holds for the asymptote of low mean rate μ → 0. That is, as long as all the steps are much slower than off-rate, their heterogeneity does not affect. In a stochastic regime, on the other hand, high mean rate μ is necessary for sufficient ϵ, and the susceptibility to rate heterogeneity is inevitable.

### Thread multiplicity offers an additional axis of performance improvement

Multi-thread scheme in a sharp firing regime has two parameters (*q*, *k_p_*) and multiple (ϵ, ϕ) trajectories are obtained for a given (*m*, *n*). We numerically pre-optimized these parameters to evaluate the best-performing receptor response (see **Methods**). The obtained receptor response is shown in **Fig. 5A**. The effective number of steps increased nearly linearly for both *n* and *m*, exhibiting near-multiplicative enhancement for the tested range, including single-step cases (**Fig. 5B**). With (*m*, *n*) = (10, 3) reflecting ten ITAM domains undergoing three-step reaction (e.g. two ITAM phosphorylation steps and Zap70 activation step), *n*_eff_ ≈ 22 readily exceeds the required level for *C* = 0.9. Multi-thread scheme in a sharp firing regime achieves optimal performance (*n*_eff_ →∞) for either large *n* or large *m* limits (**Appendix 4**). The asymptotic behavior P(*s*|*k*) ∝ *k*^−*qmn*^ for *k* ≫ *k_p_* indicates multiplicative enhancement also for deterministic limit when *q* ≈ 1 (see **Appendix 5**).

**Figure 5.**
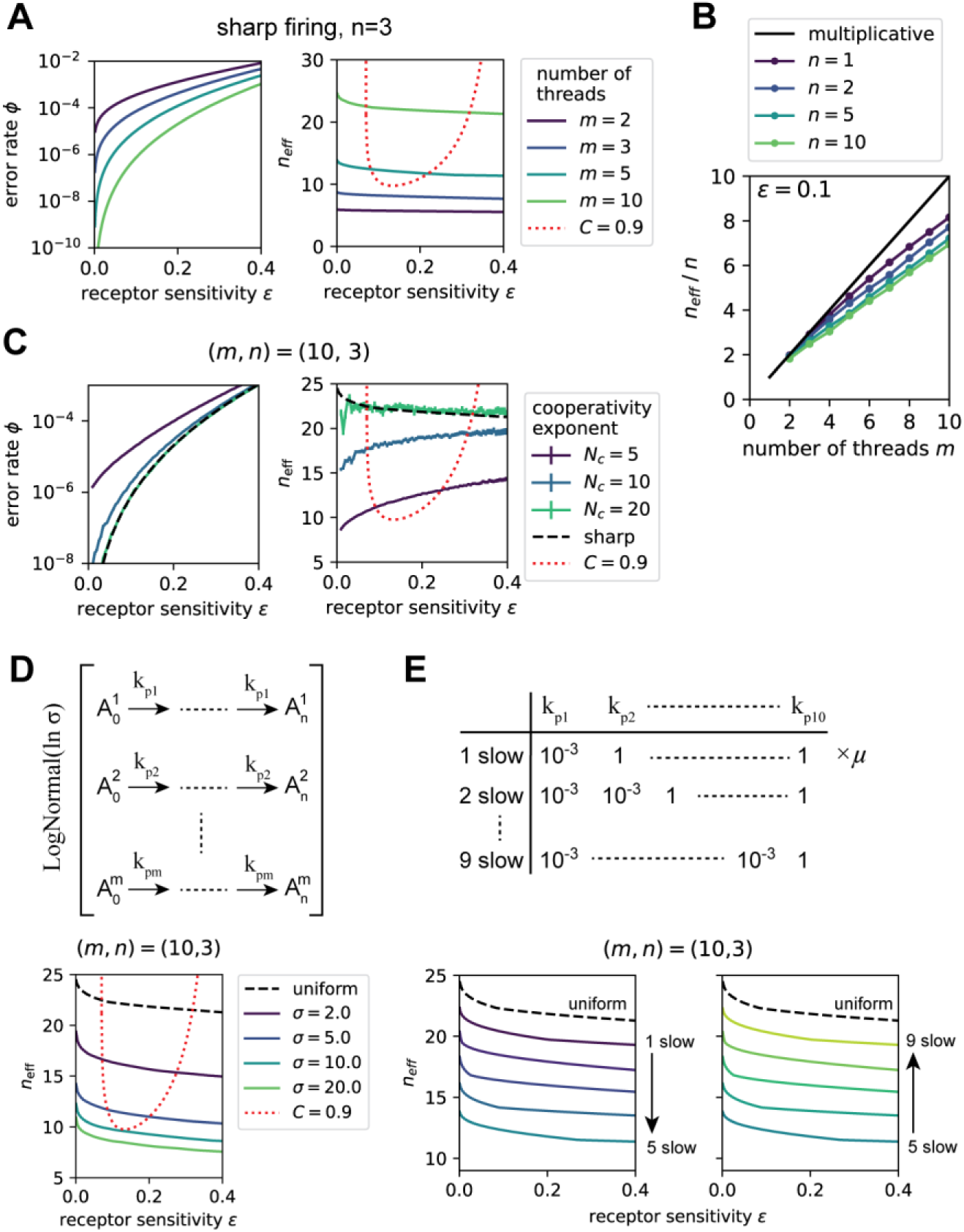
Performance of multi-thread scheme. (**A**) Receptor response trajectories (left) and effective number of steps *n*_eff_ (right) in a sharp firing regime. (**B**) *n*_eff_ sliced at ɛ = 0.1 for sharp firing with representative (*m*,*n*) pairs. The multiplicative line *n*_eff_ = *mn* is shown for comparison. (**C**) Receptor response (left) and *n*_eff_ (right) for several finite cooperative exponent *N_c_*. The curves for the corresponding model in a sharp firing regime are overlaid. (**D**) Evaluation of m-axis rate heterogeneity by stochastic sampling from log-normal distributions with different relative variation σ, in a sharp firing regime. Median values from multiple trials are shown. (**E**) Performance of multi-thread scheme in a sharp firing regime with a defined number of slow threads.

With finite cooperativity exponent *N_c_*, the expression of P(*s* = 1|*k*) includes multiple integrals and it was evaluated by Monte-Carlo method (see **Methods**). Receptor response with pre-optimized parameters is shown in **Fig. 5C**. For (*m*, *n*) = (10, 3), the effective number of steps increased with *N_c_* and it reached the sharp firing limit at *N_c_* of about 20. The performance starts to exceed *C* = 0.9 requirement at sub-optimal *N_c_* ≈ 5. Cooperative firing is essential for multi-thread scheme to enhance the performance.

There can be m-axis (inter-thread) and n-axis (intra-thread) rate heterogeneity for multi-thread scheme. We assess the m-axis heterogeneity for simplicity, because we saw that sequential heterogeneous reactions simply collapse to slowest steps. The effect of heterogeneity was evaluated by stochastic sampling in a sharp firing regime (see **Methods**). *n*_eff_ decreased as the relative variation σ increased, and the extent of degradation was similar to that of sequential scheme (**Fig. 5D** in comparison with **Fig. 4F**). The performance degradation was symmetric with respect to the number of slow/fast threads (**Fig. 5E**).

## Interpretations of the model analysis results

To achieve a good discrimination performance with the sequential scheme, all the reaction steps must have highly aligned rates. The molecular identities of the reaction steps are distinct (e.g. recruitment and activation of kinases and phosphorylation of substrates), which make them unlikely to have uniform rates as an inherent property. Therefore, the sequential scheme faces a challenge for molecular implementation to optimize the rates of many distinct reactions. Some previous studies have claimed apparently contrasting conclusions due to different model settings, and we clarify the differences in **Appendix 6**.

With the multi-thread scheme, increasing the number of threads is almost as effective as increasing the number of steps, provided that the reaction rates are aligned among threads. Contrasting the sequential scheme, simply having multiple copies of the same ITAM domains on the TCR readily achieves both the increased number of threads and the intrinsically aligned rates, since the reactions are chemically identical. The multi-thread scheme offers a new axis of performance enhancement which is potentially easier to implement, and T cell has plausibly evolved in this direction for this reason.

This conceptual advantage of multi-thread scheme holds when at least two effective steps are required, on the grounds that having two steps with aligned rates is difficult. We note that this rate alignment refers to overall reaction rate, not molecular kinetic rate constants, and is thus also highly dependent on protein expression level. As described in **Appendix 7**, we confirmed the consistency in three different model settings: (i) the performance benchmark and the parameter tuning precision may vary, (ii) the number of binding events (*N*_0_ and β) may vary to weaken the stochastic effects, (iii) the non-binary signal is continuously produced from activated receptor until unbinding, as is also common in modeling studies (5, 17).

## Re-interpretations of experimental data based on multi-thread scheme

### Relationships between T cell response and ligand off-rate

Kinetic proofreading by TCR has largely been analyzed by fitting data to sequential scheme-based models. We demonstrate here that the multi-thread scheme can consistently describe these previous data and provides new interpretations of their implications. For the first example, we analyze the affinity-potency relationship measured with a panel of peptides (16). In this study, the affinity (*K*d) and the effective concentration that induces CD69 upregulation in 15% of T cells (*P*15) were measured for various peptides. The number of proofreading steps was then estimated to be about 2.7 using a deterministic model which essentially incorporates sequential scheme (**Fig. 6A**). We rescaled *K*d and *P*15 to off-rate *k* and receptor response P(*s*|*k*) by assuming that the on-rate and the number of binding events is each uniform, and performed data fitting (**Fig. 6B, C**, see **Methods**). Fitting by sequential scheme yielded the number of steps to be 3.5. A little higher value than the original study is consistent with the introduction of stochasticity. The same data were fitted by multi-thread schemes with different number of threads, which yielded almost identical fitting curves. As the number of threads increases, the number of steps is reduced to be 1.8, 1.2, 0.9 steps for 2, 3, 4 threads, exhibiting the compensatory relationship between steps and threads. Therefore, multiple steps can be re-interpreted as multiple threads to explain experimental data with indistinguishably similar affinity-potency relationships.

**Figure 6.**
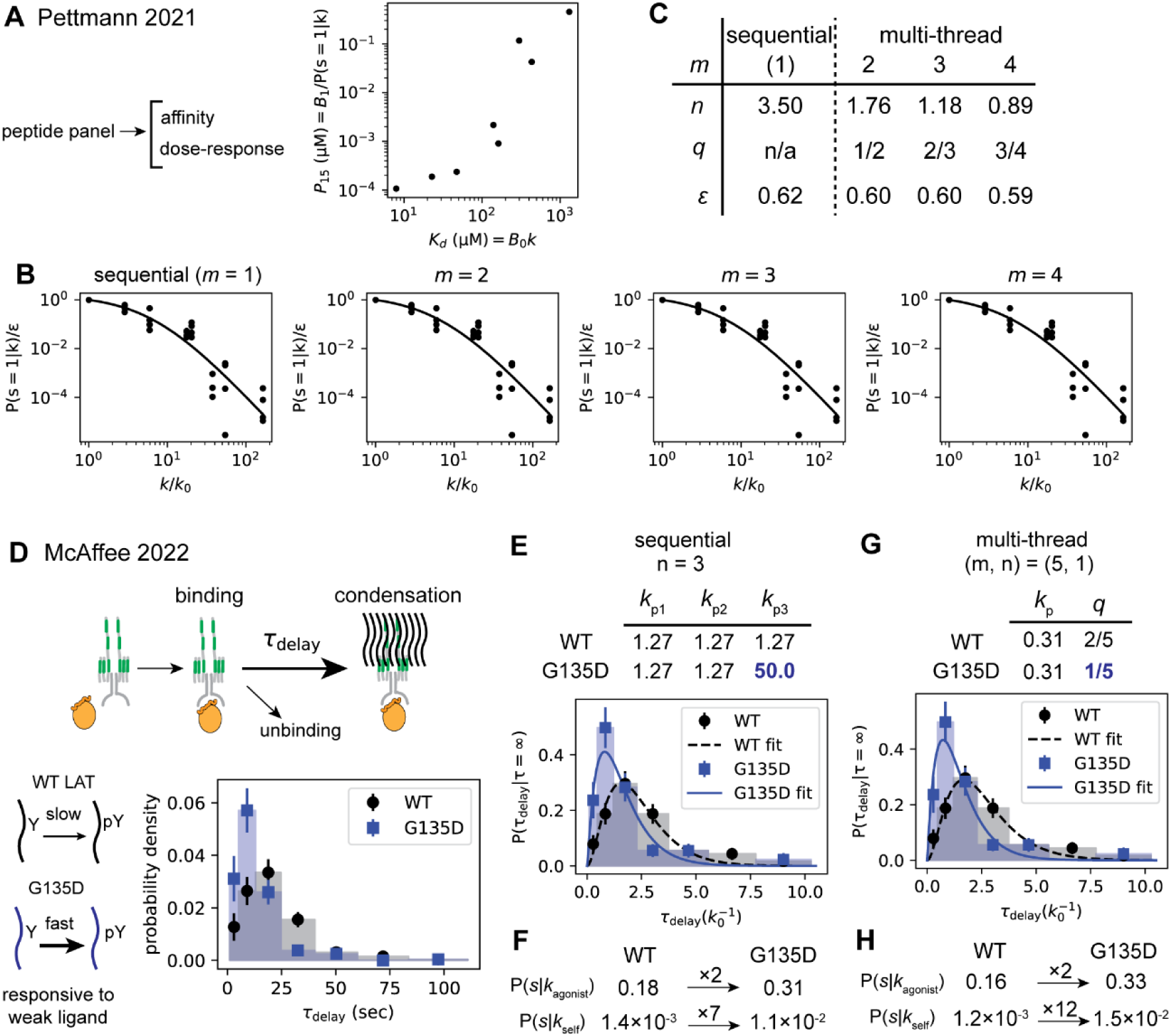
Interpretations of experimental data by sequential and multi-thread schemes. (**A**) Re-plotted affinity-potency relationships reported in the reference (16). (**B**) Rescaled and pooled data points plotted as dots (4 available experimental replicates). Fit curves by sequential scheme or multi-thread scheme with different number of threads *m* are shown. (**C**) Parameters determined from fitting. Firing threshold *q* were chosen so that *n* is minimized. (**D**) Re-plotted delay time distributions between pMHC binding and LAT condensation events reported in (38). Schematic of delay time (top), schematic of LAT G135D mutation effects (bottom left), and the distributions (bottom right) are shown. Standard errors were estimated by the square root of counts for each bin and are shown as error bars. (**E**) Internal delay time distributions after re-scaling. Data was fitted by 3-step sequential schemes. The rate of the last step for G135D was allowed to vary independently. Rates are shown in the unit of agonist off-rate *k*_0_. (**F**) Receptor response corresponding to the determined parameters for sequential scheme. (**G, H**) Internal delay time distributions fitted by 5-thread-1-step multi-thread scheme, and the resulting receptor response are shown. Firing threshold *q* was allowed to vary independently.

### LAT condensation delay time and fast-phosphorylatable LAT mutation

For the second example, we analyze our previously reported measurement of the delay time distributions between pMHC-TCR binding and LAT condensation events (38). An ensemble of delay time τ_delay_ were measured for single-molecule pMHC binding and individual LAT condensation events (**Fig. 6D**). By considering LAT condensation event as the final reaction step at which the binary signal is produced, the theoretical delay time distribution can be derived for both schemes (see **Methods**). Additionally, delay time distribution for LAT G135D mutant was also measured and found to be shorter than wildtype. This mutant exhibits faster phosphorylation at tyrosine 136, one of tyrosine residues contributing to LAT crosslinking, and it enhances the T cell response especially to weak ligands (29). For sequential scheme, the mutation effect can be interpreted as the acceleration of one of the reaction steps (30). The delay time distributions were fitted by 3-step sequential schemes, where the rate for the last step for G135D was allowed to vary, while the other steps shared the same rate (**Fig. 6E**, see **Methods**). The obtained rate for the last step for G135D was faster than the other steps, and the receptor response was enhanced preferentially for the self peptide (**Fig. 6E, F**). For multi-thread scheme, faster LAT phosphorylation corresponds to a larger nucleation rate with smaller firing threshold *q*. The delay time distributions were fitted by 5-thread-1-step multi-thread scheme in a sharp firing regime, where *q* was allowed to vary independently for each dataset (**Fig. 6G**). The firing threshold for G135D was smaller, and the resulting receptor response was also preferentially enhanced for the self peptide (**Fig. 6G, H**). Therefore, multi-thread scheme provides an alternative interpretation of the effects of G135D mutation, where the lower nucleation threshold describes both shorter delay time and enhanced response to weak ligands. We note that the number of steps and threads (3 steps or 5 threads) chosen for the fitting are just an example, and different values may fit data fairly well. Our claim here is limited to the finding that the faster proofreading step in sequential scheme can be re-interpreted as the accelerated firing step in multi-thread scheme.

### Modified multi-thread scheme recapitulates inhibitory signaling from early-state ITAMs

One way to experimentally investigate multi-thread scheme is the deletion or dysfunctional mutation of ITAM domains, which is expected to decrease the number of threads *m* while other parameters 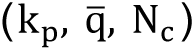 are unchanged, decreasing the sum signal produced. Previous reports are overall inconsistent and inconclusive, some reporting positive correlation of the ITAM copy number and T cell functionality (49–52), no or little correlation (53, 54), or negative correlation (36). Nonetheless, a recent counterintuitive report observing that the partial deletion of ITAM domains rather enhances the T cell response (36) motivates us to explore descriptive models. In this study, tyrosine residues of 3 ITAM domains in CD3ζ chain were mutated to phenylalanine (6F mutant), which functionally deletes 6 out of 10 ITAM domains, and it enhanced T cell response to weak ligands (**Fig. 7A**). These observations suggest inhibitory roles of ITAM domains, and the phosphatase SHP1 activity associated with partially phosphorylated ITAM was proposed as a plausible mechanism. A modified sequential scheme with inhibitory components was used to explain data (36).

**Figure 7.**
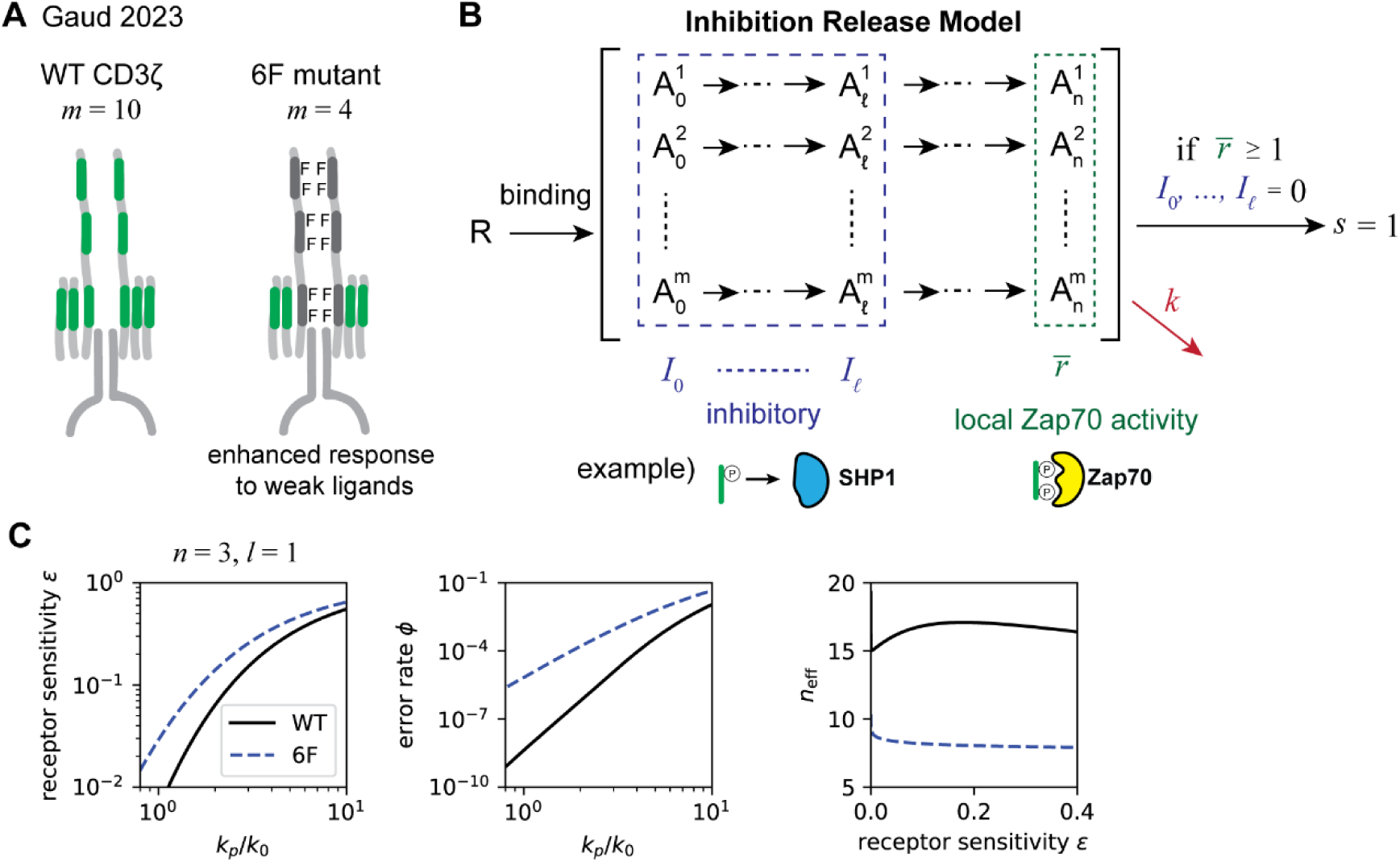
Interpretation of ITAM mutant behavior by inhibition release model. (**A**) Observations from the reference (36). TCR:CD3 complex with wild type CD3ζ corresponds to 10 threads, while the complex with 6F mutant corresponds to 4 threads in multi-thread scheme. 6F mutant exhibited enhanced response to weak ligands. (**B**) Modified multi-thread scheme with inhibition release mechanism. (**C**) (left and center) The receptor response as a function of forward reaction rate *k_p_*(in the unit of *k*_0_) for wildtype and 6F mutant with shared parameters *n* = 3 and *l* = 1. (right) Corresponding effective number of steps.

We constructed the modified multi-thread scheme incorporating inhibitory roles of ITAM domains in early states (**Fig. 7B**). In this model, while the completed threads contribute positively to the firing rate, the threads at early states up to *l*-th state negatively contribute to the firing rate. For simplicity, we suppose the strong inhibition/nucleation assumption, where one inhibitory thread completely blocks the firing, and one completed thread immediately induces firing as long as there is no inhibition (see **Methods** for details). It is notable that inhibitory signals are implemented to operate within single binding events, contrasting to feedback/feedforward models which operate among binding events (30, 33–35). Using *n* = 3 and *l* = 1 as example parameters, the calculated receptor responses for wild type and 6F mutant are shown in **Fig. 7C**. The response of 6F was enhanced slightly for agonist (ɛ increased by a factor of about 2) and drastically for self (ϕ increased by a factor of about 10^2^), recapitulating the observations. Strong inhibition/nucleation assumption additionally showed the beneficial outcome that the fine-tuning of nucleation threshold *q* is now unnecessary. While the optimization of *q* is critical for the original multi-thread scheme to achieve high performance (**Fig. S2**), inhibition-release model achieves fairly good performance without corresponding parameter tuning, which also improves just by having more threads (**Fig. 7C**, right panel). Together, multi-thread scheme was uniquely capable of incorporating inhibitory signaling as inter-thread interferences within single receptor, which recapitulated the observations.

## Discussion

Historically, kinetic proofreading by T cell receptor has been modeled by an unbranched sequence of reaction steps, which we have called the sequential scheme. In the present work, we proposed a multi-thread scheme with integrative firing step, recapitulating ITAM multiplicity and LAT condensation elements of the TCR molecular signaling mechanism. Whereas ITAM multiplicity on the TCR has been known for decades, its potential to contribute to kinetic proofreading fidelity requires an integration step, which has only recently been observed in the form of LAT condensate nucleation. Furthermore, LAT condensate nucleation model exhibits highly non-linear integration kinetics, which was revealed to be essential for the multi-thread scheme to function (see **Appendix 8** for extended discussion). Indeed, the performance enhancement in kinetic proofreading created by the LAT condensate enabling the multi-thread mechanism may well have been a driver of their evolution. We demonstrated that experimental data which has been interpreted by the sequential scheme can also be explained by the multi-thread scheme. These analyses do not provide exclusive evidence against sequential scheme, but do show that both schemes are equally consistent with data. We suggest that the multi-thread scheme more directly reflects known features of the TCR molecular signaling mechanism.

We identified the advantage of multi-thread scheme as its evolutionary accessibility, obviating the need for multi-step reactions with aligned rates. Other systems of kinetic proofreading, such as DNA replication and translation, typically consist of a single proofreading step, suggestively underpinning the evolutionary hurdle for multi-step proofreading (55, 56). As a special example, RecA protein polymerization may involve multi-step proofreading utilizing the sequential and uniform nature of elongation reactions (57, 58), which can be viewed as a strategy contrasting to TCR. TCR signaling, where elongation reactions do not participate in proofreading, might have evolved to utilize the multi-thread architecture. Notably, the roles of ITAM multiplicity are still debated (26, 36, 52, 59, 60), and the roles of LAT condensation remain largely underexplored.

## Acknowledgments

The financial support for this work was provided by NIH grant P01 AI091580 (J.T.G.), by the Novo Nordisk Foundation Challenge Programme as part of the Center for Geometrically Engineered Cellular Systems NNF17OC0028176 (J.T.G.), and by The Nakajima Foundation Scholarship (S.M.).

## Notes

### Competing Interest Statement

The authors have declared no competing interest.

