## Supporting Information for "Kinetic Proofreading through Parallel Reactions on a Single T Cell Receptor"

**This PDF file includes:**

Supporting Information Text – Methods, Appendices 1-8.
Supplementary Figures S1 to S10
SI References

### Supporting Information Text

#### Methods

Calculation of the effective number of steps

The trajectory for sequential scheme uniquely covers the entire possible range of $(\epsilon, \phi)$ space as $n$ varies (**Appendix 3**). We can define the effective number of steps $n_{\mathrm{eff}}(\epsilon, \phi) \in (0, \infty)$ as the inverse function of the trajectory for $1 > \phi > \epsilon^{\alpha-1}$. This means that the molecular system of interest is equivalent to $n_{\mathrm{eff}}$-step sequential scheme, providing an intuitive and general performance metric. $n_{\mathrm{eff}}(\epsilon, \phi)$ was numerically calculated by root-finding.

Stochastic sampling of heterogeneous forward reaction rates for sequential scheme

Receptor response for n-step sequential scheme with individual forward reaction rates $k_{pi}(i = 1, \dots, n)$ is given as $\mathrm{P}(s = 1|k) = \prod_i \{ k_{pi}/(k_{pi} + k)\}$. As numerical experiments, $\{k_{pi}\}$ were normalized as $k_{pi}/\mu = \eta_i$, which were sampled from log-normal distribution $\mathrm{P}(\eta) = \mathrm{Normal}(\ln \eta / \ln \sigma)$, so that $\{k_{pi}\}$ varies by a multiplication factor of $\sigma$ around $\mu$. $\mathrm{Normal}(\cdot)$ represents standard normal distribution. Varying $\mu$ yields a $(\epsilon, \phi)$ trajectory for a given $\{\eta_i\}$. The median $\phi$ and $n_{\mathrm{eff}}$ from 1000 sampling trials were obtained as the representative trajectories.

Receptor response calculation for multi-thread scheme in a sharp firing regime

The distribution of $\bar{\mathrm{r}}$ at time $t$ follows

$$\mathrm{P}(\bar{\mathrm{r}}, t) = \mathrm{Binomial}\left(\bar{\mathrm{r}}; m, \mathrm{CDF}_{\mathrm{Gamma}}\left(t; n, k_p\right)\right) \qquad (S1)$$

and

$$\mathrm{P}(s|k) = \int_0^\infty k e^{-k\tau} \left[ \sum_{\bar{\mathrm{r}} > \bar{q}} \mathrm{P}(\bar{\mathrm{r}}, \tau) \right] \mathrm{d}\tau \qquad (S2)$$

, where $\bar{q} = mq$ is defined so that $\bar{q} \in N$, $\mathrm{Binomial}(\cdot; n, p)$ represents binomial distribution with the number of trials $n$ and probability $p$, and $\mathrm{CDF}_{\mathrm{Gamma}}(\cdot; n, k)$ represents cumulative density function of gamma distribution with shape parameter $n$ and rate parameter $k$. This can be calculated by numerical integration. Since two parameters $(\bar{q}, k_p)$ are adjustable, multiple trajectories are obtained by changing $\bar{q}$, and the optimal performance appears as the lower envelope. Example trajectories are shown in **Fig. S2**. Numerically determined lower envelope yields the optimal paired parameters $(\bar{q}, k_p)$ (lower two panels).

Stochastic sampling of heterogeneous forward reaction rates in multi-thread scheme in a sharp firing regime

Denoting $k_{pij}$ representing the $k_p$ for j-th thread i-th step, m-axis and n-axis heterogeneity can be considered in general. Only m-axis heterogeneity is considered for simplicity. $k_{pij}$ is simplified as $k_{pj}$, and expressed as $k_{pj} = \mu_{kp}\eta_j$, where $\eta_j$ represents relative deviations. We perform stochastic sampling from log-normal distribution as $\mathrm{P}(\eta) = \mathrm{Normal}\left(\ln(\eta) / \ln(\sigma_{kp})\right)$. Then, $\mathrm{P}(\bar{\mathrm{r}}|\mathrm{k})$ is expressed with Poisson-Binomial distribution,

$$P(\bar{r}|k) = \int_0^\infty d\tau P(\tau|k) \mathrm{PoiBin}\left(\bar{r}\middle|m, \{\mathrm{CDF}_{\mathrm{Gamma}}\left(\tau\middle|n, k_{pj}\right)\}_j\right)$$
$$= \int_0^\infty d\tau' \left(k/\mu_{kp}\right) e^{-(k/\mu_{kp})\tau'} \mathrm{PoiBin}\left(\bar{r}\middle|m, \{\mathrm{CDF}_{\mathrm{Gamma}}\left(\tau'\middle|n, \eta_j\right)\}_j\right) \qquad (S3)$$

, where $\tau' = \mu_{kp}\tau$, and $\mathrm{PoiBin}\left(\cdot; m, \{p_j\}_j\right)$ is a Poison-Binomial distribution with the number of trials $m$ and a sequence of probability $\{p_j\}_{j=1,\dots,m}$. With this, $\{\eta_j\}$ was sampled and $(\epsilon, \phi)$ trajectories were calculated for each $\bar{q}$ by varying $\mu_{kp}$. Optimal $\bar{q}$ were numerically found and the corresponding $(\epsilon, \phi)$ trajectory was obtained as the optimal receptor response for each sampling trial. The median $\phi$ and $n_{\text{eff}}$ from 100 sampling trials were obtained as the representative trajectories.

Receptor response calculation for multi-thread scheme with finite cooperativity exponent

For finite $N_c$, $k_f = k_0(r/q)^{N_c}$ takes different values for each $r$. Here we denote $k_{fi} = k_0(i/\bar{q})^{N_c}$ as the firing rate for the number of completed threads $\bar{r} = i$. Suppose the sorted timing sequence of each thread completion $\{t_j\}_{j=1,\dots,m}, t_j \le t_i \text{ for } j < i$, is sampled from its probability distribution $\mathrm{P}\left(\{t_j\}\right)$. Here $\{t_j\}$ is provided for all $j = 1,2,\dots,m$, without considering the unbinding event prior to thread completion. From symmetry, $\mathrm{P}\left(\{t_j\}\right) = m! \prod_{j=1}^{m} \mathrm{PDF}_{\text{Gamma}}\left(t_j; n, k_p\right)$, where $\mathrm{PDF}_{\text{Gamma}}(\cdot; n, k)$ represents probability density function of gamma distribution with shape parameter $n$ and rate parameter $k$.

$\mathrm{P}(s = 1|k)$ for a given $\{t_j\}$ can be calculated by definition,

$$\begin{aligned}
\mathrm{P}(s = 1|k) &= \int_0^\infty k e^{-k\tau} \mathrm{P}(s = 1|\tau) d \\
&= \sum_{j=0}^{m} \int_{t_j}^{t_{j+1}} d\tau k e^{-k\tau} \{ 1 - e^{-k_{fj}(\tau - t_j)} \prod_{i=0}^{j-1} e^{-k_{fi}(t_{i+1} - t_i)} \} \\
&= 1 - \sum_{j=0}^{m} \frac{k}{k + k_{fj}} \left( e^{-(k+k_{fj})t_j} - e^{-(k+k_{fj})t_{j+1}} \right) e^{k_{fj} t_j - \sum_{i=0}^{j-1} k_{fi}(t_{i+1} - t_i)} \\
&= \sum_{j=1}^{m} \left( \frac{k}{k + k_{f(j-1)}} - \frac{k}{k + k_{fj}} \right) e^{-k t_j - \sum_{i=0}^{j-1} k_{fi}(t_{i+1} - t_i)} \qquad (S4)
\end{aligned}$$

, where $t_0 = 0$, $t_{m+1} = \infty$ were defined and $k_{f0} = 0$ was used. Integrating this over the distribution of $\{t_j\}$,

$$\mathrm{P}(s = 1|k) = \int_{0 < t_1 < \cdots < t_m < \infty} \mathrm{P}\left(\{t_j\}\right) \sum_{i=1}^{m} A_i \left( \prod_{j=1}^{m} d\, t_j \right) \qquad (S5)$$

, where

$$A_i = \left( \frac{k}{k + k_{fi-1}} - \frac{k}{k + k_{fi}} \right) e^{-\{(k + k_{fi-1}) t_i + \sum_{j=1}^{i-1} (k_{fj-1} - k_{fj}) t_j\}}$$

This integration was numerically evaluated by Monte-Carlo method, where $\{t_j\}$ was randomly sampled as a trial, and $\mathrm{P}(s = 1|k)$ was calculated. Mean and standard error of mean for $\mathrm{P}(s = 1|k)$ from multiple samples were calculated.

Since two parameters $\left(q, k_p\right)$ are continuously adjustable, lower envelope of the moving trajectory was numerically determined, representing the optimal receptor response (an example shown in **Fig. S3**). For this optimization, the $(\epsilon, \phi)$ pairs were calculated for an array of parameters. The obtained $\epsilon$ and $\phi$ as a function of $\left(q, k_p\right)$ was fit by weighted least-square bivariate spline in logarithmic space (residuals normalized by uncertainty is shown as “fit quality” in **Fig. S3B, C**), and the optimal parameter pairs were determined.

Re-analysis of affinity-potency relationships from Pettmann 2021

We re-analyzed the data reported in (1). They prepared the peptide panel for 1G4 TCR and measured the affinities and T cell CD69 responses. The affinity values $K_d$ were taken from the Source Data 2 for Figure 1 in this publication. Following the methods in this publication, the affinity value with fitted Bmax method was used for agonist peptide (9V), and the values with constrained Bmax method were used for the other peptides. The dose-response curves for 1G4 naïve T cells and CD69 response were taken from the Source Data 1 for Figure 2 in this publication. The dose-response curves were fitted by four-variable sigmoid to reproduce their analysis to obtain $P_{15}$ values. Among 6 replicates of measurements reported, 4 replicates could be used for our re-analysis, because one of the replicates had a reporting error with duplicated values, and another one had high CD69 response for all conditions and no meaningful $P_{15}$ values could be extracted. $K_d$ was assumed proportional to the unbinding rate, thus $K_d = B_0 k$ with a constant $B_0$. $P_{15}$ was assumed to be proportional to the inverse of receptor response, thus $P_{15} = B_1/\mathrm{P}(s = 1|k)$ with a constant $B_1$. The data was rescaled relative to the data point for agonist (9V peptide), which represents $K_d = B_0 k_0$ and $P_{15} = B_1/\epsilon$, and the rescaled axes $k/k_0$ and $\mathrm{P(s = 1|\,k)}/\epsilon$ were obtained. Rescaling was performed for each replicate and then 4 replicates were pooled. These rescaled values were fitted by models.

For the fitting by sequential scheme, the data was fitted by $\mathrm{P}(s = 1|k)$ expression shown in the main text, where $\mathrm{k_p}$ (or equivalently $\epsilon$) and $n$ were allowed to vary. For the fitting by multi-thread scheme in a sharp firing regime, the data was fitted by numerically calculated $\mathrm{P}(s = 1|k)$ (as described above), where $k_p$ (or equivalently $\epsilon$) and $n$ were allowed to vary. Here, $n$ was allowed to vary as any positive real number by a natural generalization of gamma distribution with non-integer $n$. For each $m$, fitting was performed for all possible $\bar{q}$, and the one that minimizes $n$ was chosen as a representative result.

Delay time distribution

Delay time $\mathrm{\tau_{delay}}$ is defined as the delay from the binding event to the final reaction step, which is the step to reach the *n*-th state for sequential scheme, and the firing step for multi-thread scheme. We define “internal delay time distribution” as the distribution of $\mathrm{\tau_{delay}}$ when unbinding event does not occur, and denote it as $\mathrm{P}(\tau_{\mathrm{delay}}|\tau = \infty)$. Then,

$$\mathrm{P}(\tau_{\mathrm{delay}}|\tau = \infty) = \frac{\delta}{\delta\tau}\mathrm{P}(s = 1|\tau) \qquad (S6)$$

When the unbinding event occurs with rate $k$, the probability of producing a signal is determined by whether the final step or the unbinding event occurs first. And the observed delay time distribution is

$$\mathrm{P}(\mathrm{\tau_{delay}}|k) = Z^{-1}\mathrm{P}(\mathrm{\tau_{delay}}|\mathrm{\tau} = \infty)\int_{\mathrm{\tau_{delay}}}^{\infty} d\mathrm{\tau}\mathrm{P}(\mathrm{\tau}|\mathrm{k}) \qquad (S7)$$

, where $Z$ is a normalization factor $\mathrm{Z} = \mathrm{P}(s = 1|k)$.

The internal delay time distribution for sequential scheme with uniform rates is

$$\mathrm{P}\left(\tau_{\mathrm{delay}}\middle|k\right) = \mathrm{PDF}_{\mathrm{Gamma}}\left(\tau_{\mathrm{delay}}; n, k_p\right) \quad (S8)$$

And that for multi-thread scheme in a sharp firing regime and uniform rates is

$$\mathrm{P}\left(\tau_{\mathrm{delay}}\middle|k\right) = (\mathrm{m} - \bar{q})\mathrm{PDF}_{\mathrm{Gamma}}\left(\tau_{\mathrm{delay}}; n, k_p\right)\times\mathrm{Binomial}\left(\bar{q}; m, \mathrm{CDF}_{\mathrm{Gamma}}\left(\tau_{\mathrm{delay}}; n, k_p\right)\right) \quad (S9)$$

, where the second term is the probability of being $\bar{r} = \bar{q}$ and the first term is the transition rate to $\bar{r} = \bar{q} + 1$.

The internal delay time distribution for the sequential scheme with heterogeneous rates were calculated numerically. The master equations for this system are expressed as

$$\frac{\partial}{\partial t}\vec{x} = M\vec{x} \quad (S10)$$

, where

$$M = \begin{bmatrix} -\left(k + k_{p1}\right) & 0 & 0 & \cdots & 0 & 0 \\ k_{p1} & -\left(k + k_{p2}\right) & 0 & \cdots & 0 & 0 \\ 0 & k_{p2} & -\left(k + k_{p3}\right) & \cdots & 0 & 0 \\ \vdots & \vdots & \vdots & \ddots & \vdots & \vdots \\ 0 & 0 & 0 & \cdots & -\left(k + k_{pn}\right) & 0 \\ 0 & 0 & 0 & \cdots & k_{pn} & 0 \end{bmatrix}$$

and

$$\vec{x} = \left(\mathrm{P}(A_0, t), \mathrm{P}(A_1, t), \dots, \mathrm{P}(A_n, t)\right)^T$$

Here $\mathrm{P}(A, t)$ is the probability of the system being state $A$ at time $t$. The internal delay time distribution was obtained by numerically solving this ODE with $k = 0$, and calculating

$$\mathrm{P}\left(\tau_{\mathrm{delay}}\middle|\tau = \infty\right) = k_{pn}\mathrm{P}\left(A_{n-1}, t = \tau_{\mathrm{delay}}\right) \quad (S11)$$

<u>Re-analysis of delay time distributions from McAffee 2022</u>

We re-analyzed our previously reported data in (2). In this report, the delay time distribution for the MCC peptide and AND TCR were measured. The distribution was also measured for the T cells transduced with LAT G135D mutant. AND TCR is a genetically modified TCR and MCC peptide-AND TCR pair exhibits stronger interaction than its native counterpart, while T102S peptide-AND TCR pair resembles the native agonist peptide binding (3). The unbinding rate for both MCC and T102S were taken from the same publication as delay time distributions (0.0227 $s^{-1}$ and 0.0926 $s^{-1}$, respectively), and T102S was treated as agonist with $k = k_0$, while MCC was treated as super agonist with $k = 0.245k_0$. The delay time distributions were represented as histograms with non-uniform bins. For each bin, the uncertainty was estimated as the square root of the counts. The distributions were converted to the internal delay time distributions and rescaled to the unit of $k_0^{-1}$ using the unbinding rate values, then fitted by models.

The experimental internal delay time distributions, $\mathrm{P}^{\mathrm{WT}}\left(\tau_{\mathrm{delay}}\middle|\tau = \infty\right)$ and $\mathrm{P}^{\mathrm{G135D}}\left(\tau_{\mathrm{delay}}\middle|\tau = \infty\right)$, were simultaneously fitted by model to minimize the summed squared residuals. For the fitting by sequential scheme, 3-step schemes with the same rate except the last step for G135D were used. The rates for WT are all $k_p$ and the rates for G135D are $k_p$ for the first and second steps and $k_p$' for the last step. The theoretical internal delay time distributions were calculated as described above and fitted to the experimental distributions. Constant scaling factors $A^{\mathrm{WT}}$ and $A^{\mathrm{G135D}}$ were also used for the fitting as

$$\mathrm{P}_{\mathrm{experimental}}^{\mathrm{WT}}\left(\tau_{\mathrm{delay}}\middle|\tau = \infty\right) = A^{\mathrm{WT}}\mathrm{P}_{\mathrm{theoretical}}^{\mathrm{WT}}\left(\tau_{\mathrm{delay}}\middle|\tau = \infty\right) \quad (S12)$$

$$\mathrm{P}_{\mathrm{experimental}}^{\mathrm{G135D}}\left(\tau_{\mathrm{delay}}\middle|\tau = \infty\right) = A^{\mathrm{G135D}}\mathrm{P}_{\mathrm{theoretical}}^{\mathrm{G135D}}\left(\tau_{\mathrm{delay}}\middle|\tau = \infty\right) \quad (S13)$$

, because the experimental distributions have large uncertainties.

For the fitting by multi-thread scheme in a sharp firing regime, 5-thread-1-step schemes were used. The rates are uniform for both distributions, but $\bar{q}$ values (or equivalently $q$ values) were allowed to vary independently. Constant scaling factors were also used for the fitting. Since $\bar{q}$ is discrete, all possible $\bar{q}$ combinations were tested, and the parameters that minimize the residuals were found.

<u>Receptor response for inhibition release model</u>

For each thread in multi-thread scheme, the probability of being state $A_i$ at time τ is

$$P_i = \frac{\left(k_p\tau\right)^i}{i!}exp\left(-k_p\tau\right) \ \ (i = 0,1,\dots,n-1) \quad (S14)$$

$$P_n = \mathrm{CDF}_{\mathrm{Gamma}}\left(\tau\middle|n,k_p\right) \quad (S15)$$

Define $I_i$ as the count of the threads at state $A_i$. The probability of satisfying the condition $I_0, I_1, \dots, I_l = 0$ and $\bar{r} > \bar{q}$ is

$$\mathrm{P}(s = 1|\tau) = \sum_{\bar{r}=\bar{q}+1}^{m} \mathrm{Binomial}(\bar{r}|m,P_n) \times \left(\frac{1-(P_0+P_1+\cdots+P_l)}{1-P_n}\right)^{(m-\bar{r})} \quad (S16)$$

The receptor response $\mathrm{P}(s = 1|k)$ was obtained by numerical integration using this expression.

**Appendix 1. Master Equations and Solution for LAT Condensate Nucleation**

The number of phosphorylated LAT tyrosine residues $N$ increases by the rate $k^{+}r$, where $\mathrm{r} \in [0,1]$ represents the local Zap70 level (**Fig. 4B**). $N$ also decreases by the rate $k^{-}N$ by phosphatase activity. Once $N$ reaches the critical number $N_c$, nucleation occurs. Thus $\mathrm{P(N,t)}$ is absorbed at $N_c$, and the momentary probability of nucleation, which we call the firing rate $k_f$, is represented by the first passage time $\tau_{\mathrm{FPT}}$.

$$\frac{d}{dt}\mathrm{P}(N,t) = k^{-}\{N_{ss}P(N-1,t) - (N_{ss}+N)\mathrm{P}(N,t) + (N+1)\mathrm{P}(N+1,t)\} \qquad (S17)$$

$$\mathrm{P}(\tau_{\mathrm{FPT}}) = k^{-}N_{ss}\mathrm{P}(N_c - 1, t = \tau_{\mathrm{FPT}}) \qquad (S18)$$

where $N_{ss} = k^{+}r/k^{-}$ represents steady-state average $N$, and $\mathrm{P}(-1,t) = \mathrm{P}(N_c,t) = 0$. The firing rate follows $k_f(t) = \mathrm{P}(\tau_{\mathrm{FPT}} = t)/\mathrm{P}(N < N_c, t)$. We consider the regime of $N_{ss} \ll N_c$ for all $r$, where the firing is always driven by fluctuation rather than build-up of $N$, and we also assume $N_c \geq 2$ so that multiple phosphorylation events are necessary.

The steady-state solution for non-absorbing system (remove the absorbing boundary $\mathrm{P}(N_c,t) = 0$ and let $N$ any non-negative integer) is easily obtained as Poisson distribution:

$$P_{ss}(N) = \frac{e^{-N_{ss}}N_{ss}^{N}}{N!} \qquad (S19)$$

Additionally, when $N$ is limited below $N_c$ (so that the state $N = N_c - 1$ can only transition to the state $N = N_c - 2$), the steady-state distribution is also equal to this with a normalization factor $\sum_N \mathrm{P}(N)$. Since $N_{ss} \ll N_c$, this normalization factor is close to 1 and negligible in the following.

If firing is much slower than fluctuation, and if the introduction of absorbing boundary at $N_c$ only gives a small perturbation, we can use the pre-equilibrium approximation. Indeed, the firing transition rate from state $N = N_c - 1$ to $N = N_c$ is given by $k^{-}N_{ss}\mathrm{P}(N_c - 1, t)$, while the dephosphorylation transition from state $N_c - 1$ to $N_c - 2$ rate is given by $k^{-}(N_c - 1)\mathrm{P}(N_c - 1, t)$, which shows that firing is much slower than dephosphorylation because $N_{ss} \ll N_c$. Thus, the introduction of firing causes only small local perturbation on $\mathrm{P}(N_c - 1, t)$. Also, the time scale of global equilibration is given by a rate $k^{-}$, because the process is essentially the counting of particles with creation rate of $k^{-}N_{ss}$ and annihilation rate of $k^{-}$. And we can later confirm that this is also much faster than the firing rate, justifying the pre-equilibrium approximation.

Under the approximation that the states below $N_c$ is pre-equilibrated, the absorption $\mathrm{P}(\tau_{\mathrm{FPT}})$ follows first-order kinetics and the firing rate is a constant,

$$k_f = k^{-}\frac{e^{-N_{ss}}N_{ss}^{N_c}}{(N_c - 1)!} \qquad (S20)$$

This is much slower than the equilibration time scale, validating the pre-equilibration as mentioned above:

$$\mathrm{k}^{-}\frac{\mathrm{e}^{-\mathrm{N_{ss}}}\mathrm{N_{ss}^{N_c}}}{(\mathrm{N_c} - 1)!} < \mathrm{k}^{-}\big(2\pi(\mathrm{N_c} - 1)\big)^{-1/2} N_{ss}\mathrm{e}^{-\mathrm{N_{ss}}}\left(\frac{eN_{ss}}{N_c - 1}\right)^{\mathrm{N_c}-1} \ll k^{-}$$

This analytical solution was also validated by comparison with numerical solution of master equations (**Fig. S4**). Since $N_{ss} \propto r$, we obtained the scaling of firing rate $k_f$ over Zap70 level $r$.

Writing $N_{ss} = rN'$, we get $k_f \propto e^{-rN'} r^{N_c}$. And $\frac{d\ln k_f}{d\ln r} = N_c - rN'$ is obtained. Since $N' \ll N_c$ and $r \in [0,1]$, $k_f \propto r^{N_c}$ is obtained as the approximate scaling.

**Appendix 2. Supplementary rationale for model formulation**

We modeled the APC presenting only agonist pMHC instead of the mixture of self and agonist for the following reason. With the model in which the mixture is presented ($N_0$ agonist and $\beta N_0$ self binding events), the antigen discrimination becomes perfect regardless of the receptor selectivity in the deterministic limit (large $N_0$ limit). Each type of ligand produces a fixed amount of sum signal in this limit. Even if self pMHC produces more signal than agonist, antigen discrimination is perfect because agonist plus self deterministically produces larger signal than only self. This anomalous behavior appears because we modeled the number of binding events $N$ as Poisson distribution, assuming a fixed ligand abundance and a fixed sensing time, and $N$ will be deterministic when sufficiently large. Realistically, the number of binding events $N$ should have additional variations, because the ligand abundance, receptor abundance, sensing time and so on vary among different APC-T cell encounter events. More accurate modeling of $N$ would be achieved by adding a noise component with $O(N)$ scaling, which has been done in a previous study (4), but it sacrifices the simplicity of the model. Instead, we adopted the comparison of agonist-only and self-only, where agonist pMHC must produce larger signal than self pMHC in order to achieve antigen discrimination. This roughly corresponds to the noise component of $\pm N$, scaling as $O(N)$, approximating the noise component discussed above.

We modeled the number of binding events to be proportional to the abundance of peptides. Experimentally, the fraction of bound pMHC was reported low from 20% to 40% for MCC-T102S peptide and AND-TCR pair (representing the affinity of agonist peptide) (3). The receptor occupancy is also low because TCR and self pMHC are similarly abundant (about $100-1000\ molecules\ \mu\mathrm{m}^{-2}$) (5, 6) with low fraction bound ($< 4\%$ inferred from agonist). In this regime where most receptor and ligand molecules are unbound, the frequency of binding events is proportional to the abundances of two species.

**Appendix 3. Sequential scheme covers the entire possible range of receptor response.**

Consider the $(\epsilon, \phi)$ trajectory of sequential scheme with generalized $n \in \mathbb{R}^+$,

$$\phi = \epsilon^{-1}\left(\alpha\left(\epsilon^{-\frac{1}{n}} - 1\right) + 1\right)^{-n}$$

Asymptotic trajectory at $n \to 0$ converges to $\phi = 1$. $\phi$ strictly decreases with $n$ for fixed $\epsilon$ (**Appendix 3-a**). Asymptotic behavior at $n \to \infty$ converges to sharp threshold of $\tau$ (7),

$$\mathrm{P}(s = 1|\tau) = \begin{cases} 1\ (\text{for } \tau > \tau_c) \\ 0\ (\text{for } \tau \le \tau_c) \end{cases} \qquad (S21)$$

And the trajectory converges to

$$\phi = \epsilon^{\alpha - 1} \qquad (S22)$$

where $\tau_c$ acts as a parameter for this trajectory. This receptor response is optimal: any molecular system cannot achieve a lower error rate than this sharp threshold (**Appendix 3-b**). Thus, the trajectory for sequential scheme uniquely covers the entire possible range of $(\epsilon, \phi)$ space as $n$ varies.

**Appendix 3-a. Proof that the sequential scheme's trajectory is monotonic**

We prove that

$$\phi = \epsilon^{-1}\left(\alpha\left(\epsilon^{-\frac{1}{n}} - 1\right) + 1\right)^{-n}$$

strictly decreases with $n$ for $n \in (0, \infty)$, $\alpha > 1$, $\epsilon \in (0,1)$.

*Proof.*

We prove that the following is negative.

$$\frac{\partial}{\partial n}\ln\phi = -\ln(x+1) - \frac{(\ln\epsilon)(x+\alpha)}{n(x+1)}$$
$$= (x+1)^{-1}\left[(x+\alpha)\ln\left(\frac{x+\alpha}{\alpha}\right) - (x+1)\,ln\left(\frac{(x+1)}{1}\right)\right]$$

, where $x = \alpha\left(\epsilon^{-\frac{1}{n}} - 1\right)$, and $x \in (0, \infty)$. Since $\alpha > 1$, the negativity of this is sufficiently proven by proving that $f(y) = y\ln\left(\frac{y}{y-x}\right)$ strictly decreases with $y$ for $y \geq 1 + x$.

$$\frac{df}{dy} = \ln\left(1 + \frac{x}{y-x}\right) - \frac{x}{y-x} = \ln(1+z) - z$$

where $z = \frac{x}{y-x}$ and $z \in (0, \infty)$. This is negative for $z > 0$, which gives the proof. □

**Appendix 3-b. Proof that sharp threshold is optimal**

For $s \in \{0,1\}$, sharp threshold

$$\mathrm{P}(s=1|\tau) = \begin{cases} 1\ (\tau > \tau_c) \\ 0\ (\tau \leq \tau_c) \end{cases}$$

, where $\tau_c \in (0, \infty)$ is a parameter, is optimal. Here, being optimal means that this $\mathrm{P}(s=1|\tau)$ as a function of $\tau$ minimizes $\phi = \mathrm{P}(s=1|k=\alpha k_0)/\mathrm{P}(s=1|k=k_0)$ for any $\epsilon = \mathrm{P}(s=1|k=k_0)$. Further, optimal $\mathrm{P}(s=1|\tau)$ is uniquely determined almost everywhere.

The proof is given by the lemma below, using the fact that

$$\frac{P(\tau|k=\alpha k_0)}{P(\tau|k=k_0)} = \alpha e^{-(\alpha-1)k_0\tau}$$

strictly decreases and satisfies the condition for the lemma. Notions like “almost everywhere” are included for completeness, but it does not affect our discussion which focuses only on "well-behaved" probability distributions.

*Lemma.*

Let $P(\tau)$ a function $P: (0, \infty) \to [0,1]$. Let $f(\tau)$ and $g(\tau)$ functions with the same domain, which satisfies (i) $f(\tau) > 0$ and $g(\tau) > 0$, (ii) $\int_0^\infty f(\tau)d\tau = 1$ and $\int_0^\infty g(\tau)d\tau = 1$, and (iii) $h(\tau) = \frac{g(\tau)}{f(\tau)}$ is strictly decreasing. Then, for given $a \in (0,1)$, the function $P$ that satisfies $\int_0^\infty f(\tau)P(\tau)d\tau = a$ and minimizes $b = \int_0^\infty g(\tau)P(\tau)d\tau / \int_0^\infty f(\tau)P(\tau)d$ is uniquely determined as

$$P(\tau) = \begin{cases} 1\ (\tau > \tau_c) \\ 0\ (\tau \le \tau_c) \end{cases}$$

almost everywhere, where $\tau_c \in (0, \infty)$ satisfies $a = \int_{\tau_c}^{\infty} f(\tau) d\tau$.

*Proof.*

$\tau_c$ is unique because $\int_{\tau_c}^{\infty} f(\tau) d\tau$ strictly decreases with $\tau_c$. Let

$$Q(\tau) = \begin{cases} 1\ (\tau > \tau_c) \\ 0\ (\tau \le \tau_c) \end{cases}$$

and express $P(\tau)$ as

$$P(\tau) = \begin{cases} Q(\tau) + R_1(\tau)(\tau \le \tau_c) \\ Q(\tau) - R_2(\tau)(\tau > \tau_c) \end{cases}$$

Then $R_1(\tau) \ge 0$ and $R_2(\tau) \ge 0$. It has to satisfy

$$\mathrm{a} = \int_{\tau_c}^{\infty} f(\tau) d\tau + \int_0^{\tau_c} f(\tau) R_1(\tau) d\tau - \int_{\tau_c}^{\infty} f(\tau) R_2(\tau) d\tau$$

then

$$\int_0^{\tau_c} f(\tau) R_1(\tau) d\tau = \int_{\tau_c}^{\infty} f(\tau) R_2(\tau) d\tau \ge 0$$

Thus,

$$\begin{aligned} b &= \frac{\int_0^{\infty} g(\tau) Q(\tau) d\tau + \int_0^{\tau_c} g(\tau) R_1(\tau) d\tau - \int_{\tau_c}^{\infty} g(\tau) R_2(\tau) d\tau}{a} \\ &= \frac{\int_0^{\infty} g(\tau) Q(\tau) d\tau + \int_0^{\tau_c} f(\tau) h(\tau) R_1(\tau) d\tau - \int_{\tau_c}^{\infty} f(\tau) h(\tau) R_2(\tau) d\tau}{a} \\ &\ge \frac{\int_0^{\infty} g(\tau) Q(\tau) d\tau + h(\tau_c) \int_0^{\tau_c} f(\tau) R_1(\tau) d\tau - h(\tau_c) \int_{\tau_c}^{\infty} f(\tau) R_2(\tau) d\tau}{a} \\ &= \frac{\int_0^{\infty} g(\tau) Q(\tau) d\tau}{a} \end{aligned}$$

Here the condition (iii) was used. Equality holds only when $R_1 = 0$ and $R_2 = 0$ almost everywhere. This is equivalent to the proof. □

**Appendix 4. Multi-thread scheme in a sharp firing regime reaches optimum with large numbers of steps or threads.**

Here we show that multi-thread scheme in a sharp firing regime also reaches the optimal sharp dwell time threshold model for either $n \to \infty$ or $m \to \infty$.

For $n \to \infty$, each thread completion follows sharp threshold of dwell time, and $r$ as a function of $\tau$ also follows sharp threshold. Therefore, the overall response $\mathrm{P}(s|\tau)$ also follows sharp threshold with $q$ appropriately set between 0 and 1.

For $m \to \infty$, $r$ as a function of $\tau$ becomes deterministic, because the distribution of $r$ following

$$\mathrm{P}(\mathrm{r}, t) = m^{-1} \mathrm{Binomial}\left(\mathrm{mr}; m, \mathrm{CDF}_{\mathrm{Gamma}}\left(t; n, k_p\right)\right)$$

becomes infinitely sharp and deterministic as

$$\mathrm{r(t)} = \mathrm{CDF}_{\mathrm{Gamma}}\left(t; n, k_p\right)$$

For a sharp firing regime, firing is determined by the threshold $r > q$, and it occurs at the fixed timing satisfying

$$\mathrm{CDF}_{\mathrm{Gamma}}(t;n,k_p)=q$$

**Appendix 5. Analytical expression of receptor response for multi-thread scheme in a sharp firing regime**

Here we derive the analytical expression of $\mathrm{P}(s|k)$ for multi-thread scheme in a sharp firing regime. We will obtain the following expression.

$$\mathrm{P}(s|k)=\sum_{J'=(\bar{q}+1)}^{m}\mathrm{C}_{J'}^{m}\left[\sum_{i=0}^{J'-(\bar{q}+1)}(-1)^{i}\,\mathrm{C}_{i}^{J'}\right]F\left(J'\middle|k,k_p,n\right)\qquad(S23)$$

, where

$$F\left(J'\middle|k,k_p,n\right)=J'!\left[\prod_{i=1}^{J'}\left(\frac{k_p}{k+ik_p}\right)^{n}\right]\sum_{l_1=0}^{(n-1)}\sum_{l_2=0}^{l_1+(n-1)}\cdots\sum_{l_{J'-1}=0}^{l_{J'-2}+(n-1)}\left[\prod_{j=1}^{J'-1}\mathrm{C}_{(\mathrm{n}\text{-}1)}^{(\mathrm{l_j}+\mathrm{n}\text{-}1)}\left(\frac{k+jk_p}{k+(j+1)k_p}\right)^{l_j}\right]$$

and $\mathrm{C}_b^a=\frac{a!}{b!(a-b)!}$ means combination.

Firstly, define

$$A(a,i,j)=\int_0^{\infty}e^{-at}t^{i}\left(\gamma(n,k_pt)\right)^{j}dt$$

for non-negative integers $i,j$ and $a\in(0,\infty)$, where $\gamma(\cdot,\cdot)$ is the lower incomplete gamma function. Note that $n\in\mathbb{N}$ and $k_p\in(0,\infty)$. By partial integration,

$$\mathrm{A}(a,i(>0),j=0)=a^{-(i+1)}\Gamma(i+1)$$
$$\mathrm{A}\big(a,i=0,j(>0)\big)=\left(ja^{-1}k_p^{n}\right)\mathrm{A}\left(a+k_p,n-1,j-1\right)$$
$$\mathrm{A}\big(a,i(>0),j(>0)\big)=(ia^{-1})\mathrm{A}(a,i-1,j)+\left(ja^{-1}k_p^{n}\right)\mathrm{A}\left(a+k_p,i+(n-1),j-1\right)$$

, where $\Gamma(\cdot)$ is the gamma function. Then recursively,

$$\begin{aligned}A(a,i,j)&=(ia^{-1})A(a,i-1,j)+\left(ja^{-1}k_p^{n}\right)A\left(a+k_p,i+(n-1),j-1\right)\\&=(ia^{-1})\left((i-1)a^{-1}\right)A(a,i-2,j)\\&\quad+(ia^{-1})\left(ja^{-1}k_p^{n}\right)A\left(a+k_p,(i-1)+(n-1),j-1\right)\\&\quad+\left(ja^{-1}k_p^{n}\right)A\left(a+k_p,i+(n-1),j-1\right)\\&=\cdots=ja^{-(i+1)}k_p^{n}i!\sum_{l=0}^{i}(l!)^{-1}a^{l}A\left(a+k_p,l+n-1,j-1\right)\end{aligned}$$

Further recursively,

$$\begin{aligned}A(k,0,m)&=\left(mk^{-1}k_p^{n}\right)A\left(k+k_p,n-1,m-1\right)\\&=k^{-1}m(m-1)(n-1)!\,k_p^{n}\left(\frac{k_p}{k+k_p}\right)^{n}\sum_{l_1=0}^{(n-1)}(l_1!)^{-1}\left(k+k_p\right)^{l_1}A\left(k+2k_p,l_1+(n-1),m-2\right)\end{aligned}$$

$$=\cdots=k^{-1}m!\left((n-1)!\right)^{m}\left[\prod_{i=1}^{m}\left(\frac{k_p}{k+ik_p}\right)^{n}\right]\sum_{l_1=0}^{(n-1)}\sum_{l_2=0}^{l_1+(n-1)}\cdots\sum_{l_{m-1}=0}^{l_{m-2}+(n-1)}\left[\prod_{j=1}^{m-1}\mathrm{C}_{(\mathrm{n}\text{-}1)}^{(\mathrm{l_j}+\mathrm{n}\text{-}1)}\left(\frac{k+jk_p}{k+(j+1)k_p}\right)^{l_j}\right]$$

Next, define

$$F(m|k,k_p,n) = \int_0^\infty ke^{-kt}\left(\mathrm{CDF}_{\mathrm{Gamma}}(t;k_p,n)\right)^m dt$$

Then,

$$F(m|k,k_p,n) = kA(k,0,m)\Gamma(n)^{-m}$$

$$= m!\left[\prod_{i=1}^{m}\left(\frac{k_p}{k+ik_p}\right)^n\right]\sum_{l_1=0}^{(n-1)}\sum_{l_2=0}^{l_1+(n-1)}\cdots\sum_{l_{m-1}=0}^{l_{m-2}+(n-1)}\left[\prod_{j=1}^{m-1}\mathrm{C}_{(\mathrm{n}-1)}^{(\mathrm{l_j}+\mathrm{n}-1)}\left(\frac{k+jk_p}{k+(j+1)k_p}\right)^{l_j}\right]$$

Lastly,

$$P(s|k) = \sum_{J=(\bar{q}+1)}^{m}\int_0^\infty ke^{-kt}\mathrm{Binomial}(J;m,\mathrm{CDF}_{\mathrm{Gamma}}(t;k_p,n))dt$$

$$= \sum_{J=(\bar{q}+1)}^{m}\mathrm{C}_{\mathrm{J}}^{\mathrm{m}}\int_0^\infty ke^{-kt}\left(\mathrm{CDF}_{\mathrm{Gamma}}(t;k_p,n)\right)^J\left(1-\mathrm{CDF}_{\mathrm{Gamma}}(t;k_p,n)\right)^{m-J}$$

$$= \sum_{J=(\bar{q}+1)}^{m}\mathrm{C}_{\mathrm{J}}^{\mathrm{m}}\sum_{i=0}^{m-J}(-1)^i\mathrm{C}_{\mathrm{i}}^{(\mathrm{m}-\mathrm{J})}F(i+J|k,k_p,n)$$

$$= \sum_{J'=(\bar{q}+1)}^{m}\mathrm{C}_{\mathrm{J'}}^{\mathrm{m}}\left[\sum_{i=0}^{J'-(\bar{q}+1)}(-1)^i\mathrm{C}_{\mathrm{i}}^{\mathrm{J'}}\right]F(J'|k,k_p,n)$$

, where the substitution $J' = J + i$ was used.

**Appendix 6. Comparisons with relevant studies of kinetic proofreading with sequential scheme**

While we saw that larger number of steps provide better performance, several studies state apparently contrasting conclusions regarding the effectiveness of sequential scheme in a stochastic regime. Although it is common to analyze the chemical master equations instead of explicit binding events, in many cases binding events are treated as independent and identical, and these studies can be re-interpreted in the framework of explicit binding events. In this re-interpretation, the dependence of the number of binding events on several parameters needs careful attention. Specifically, the system's behavior changes depending on (i) the abundance of peptides, (ii) receptor occupancy, and (iii) ligand sequestration/rebinding (**Fig. S5**). While we assumed rare agonist, low receptor occupancy and low ligand sequestration based on experimental observations, different studies adopt different assumptions, which affect the conclusions. In addition to these factors, the explored parameter space for sequential scheme also varies among studies. While we analyzed the full space of $n$ and $k_p$, it is often the case to vary only one of them, which only shows a slice of possible receptor response.

Kirby and Zilman analyzed stochastic signal from a single receptor with variable $n$ and $k/k_p$ (ratio of forward reaction rate and off-rate), and concluded that increasing the number of steps generally degrades the antigen discrimination performance (8). They varied the ligand off-rate instead of forward reaction rate (relative to the total time), and the parameter settings were such that the receptor occupancy transitions from low regime to high regime. This transition saturates the receptor output by decreasing the number of binding events as off-rate decreases, which may lead to their distinct conclusion from ours. Additionally, they compared two ligands of the same abundance, which also shifts the performance compared to the present work. In response to this work, Xiao and Galstyan showed that the performance generally improves with more steps, when the signal integration time is allowed to vary so that the sum signal produced from agonist remains constant (9). In our model, this operation is equivalent to varying $N_0$ so that $\epsilon N_0$ is constant, while varying $n$ with fixed $k_p$. Then the performance is solely determined by $\phi$, which surely decreases with $n$. Thus, their conclusion can be consistently seen in our model, though it is not our main conclusion. Li and Chou considered the sharp threshold on dwell time (large $n$ limit) and analyzed the dynamics of response from a single receptor (10). They found that the benefits of multi-step schemes depend on the post-processing of dynamic receptor output. Since we consider the situation where a receptor binds to mostly zero or one ligand, their insights do not directly relate to our observation. Morgan and Lindsay analyzed the extreme case where T cell detects a single productive binding event, thereby the first passage time determines the T cell activation (11). They essentially only considered the fixed $k_p$ (though they varied the entire time scale, it does not change $k_p$ relative to off-rate), and varied the T cell-APC contact duration, which effectively varies the number of binding events. They found that the large number of proofreading steps is beneficial in this extreme case. Although their model formulation significantly differs, their conclusion essentially aligns with our observation that large $n$ is beneficial. Roughly, their model corresponds to the situation where the $S_c$ (threshold on sum signal $S$) is between 0 and 1, and $N_0$ can be adjusted. Since $\epsilon N_0$ can be adjusted to be larger than 1, the discrimination performance solely depends on $\phi$. Following the same logic as the analysis of Xiao and Galstyan, the observation that large $n$ is beneficial can be reproduced in our model.

**Appendix 7. Robustness of the multi-thread scheme's benefit in different settings**

In the main text, the performance benchmark $C = 0.9$ was arbitrarily set, and we also did not consider how precise the parameters need be. To obtain more comprehensive understanding, we analyzed the parameter-performance relationship by evaluating the performance $C$ for varied forward reaction rate $k_p$ for sequential scheme (**Fig. S6A**). $k_p$ has an optimal value ($k_{p,optimal}$) and $C$ exhibits a peak around it. As the number of steps $n$ increases, the peak gets higher and wider, showing that larger $n$ is generally preferable. Similarly, increasing $n_{eff}$ by increasing $n$ and $m$ resulted in higher $C$ over larger regime of parameters $(k_p, q)$ for multi-thread scheme in a sharp firing regime (**Fig. S6B**). Additionally, under any criteria for performance and parameter precision, the effective number of steps must be larger than 3 due to a bound $\beta^{-1} > \alpha^{-n}$, otherwise self always produces more sum signal than agonist. Therefore, more steps are better and no less than three steps are necessary, regardless of how we define the benchmark. This confirms the multi-thread scheme's benefit as discussed in the main text.

Although we used experimental values for the numbers of binding events, these values may depend on the environment. More agonist binding events (larger $N_0$) decrease the stochastic noise, and smaller abundance ratio $\beta$ tolerates larger error rate. To characterize how robust the multi-thread scheme's advantage is to these variations, parameter-performance relationships were evaluated for 10-times larger $N_0$ and/or 10-fold smaller $\beta$ (**Figs. S7** and **S8**). To achieve same performance $C$, smaller $n_{eff}$ was required for larger $N_0$ or smaller $\beta$. With large $n_{eff}$, both peak height and width saturated, suggesting that $n_{eff}$ beyond this point is no longer beneficial. For the condition with larger $N_0$ and smaller β, saturation occurs at about 4 steps (**Fig. S7C**). This is still more than two steps, but it might raise a concern that sequential scheme might be no longer challenging and multi-thread scheme might be no longer beneficial. To confirm that this is not the case, we performed a more detailed characterization for this condition (**Fig S9**). Noting that $N_0$ or β does not affect the trajectory of $(\epsilon, \phi)$ corresponding to a given molecular system, only the discrimination performance $C$ as a function of $(\epsilon, \phi)$ was re-evaluated. In this condition, the required $\epsilon$ is low and $\phi$ is high, and the required $n_{eff}$ is low at about 3 for $C = 0.9$ (**Fig. S9A**). The optimal $\epsilon$ is smaller at about 0.05, reflecting smaller stochastic noise. Even though the system is closer to a deterministic limit, rate heterogeneity still significantly degrades the performance to a similar degree for two schemes (**Fig. S9B, C**). Therefore, even for this condition, more than two steps with uniform rates are required, which confirms that multi-thread scheme is robustly beneficial.

We primarily focused on the binary signal $s$ produced from binding events, reflecting the experimental observation of discrete LAT condensation. On the other hand, it is also common for modeling studies to have a continuous signal produced after kinetic proofreading, and this would be interesting to consider. In a common continuous model with sequential scheme, once the ligated receptor reaches the final state, it keeps converting a substrate to a product until the ligand unbinds (**Fig. S10A**). The amount of the product is treated as the signal. Suppose that the final state immediately produces a binary signal $s_1$ and the continuous signal $s_2$ at the same time. Let $\tau_f$ the timing when the final state is reached after the binding event. By normalization, we can define $s_2 = \tau - \tau_f$. We also define the summed signal $S_1$ and $S_2$ for $s_1$ and $s_2$, respectively. We can also formulate the corresponding continuous model for multi-thread scheme, where the ligated receptor transition to $s_1 = 1$ state at the firing, then $s_2$ is produced until the unbinding event. By this formulation, the comparison of $S_1$ and $S_2$ tells how continuous models compare with binary models. Here, $S_2$ distribution is merely a conversion of $S_1$ distribution, because firstly

$$\mathrm{P}(s_2|k) = \int_0^\infty d\tau_f\, \mathrm{P}(\tau_f)\mathrm{P}(\tau = \tau_f + s_2|k) = \int_0^\infty d\tau_f \mathrm{P}(\tau_f) k\, e^{-k(\tau_f + s_2)} = e^{-ks_2}\mathrm{P}(s_1 = 1|k)$$

Then,

$$\mathrm{P}(s_2|s_1 = 1) = \mathrm{P}(s_2|k)/\mathrm{P}(s_1 = 1|k) = e^{-ks_2}$$

, showing that $s_2$ per fired receptor follows exponential distribution. Therefore,

$$\mathrm{P}(S_2) = \begin{cases} \mathrm{P}(S_1 = 0) \quad (S_2 = 0) \\ \sum_{S_1=1}^{\infty} \mathrm{P}(S_1)\mathrm{PDF}_{\mathrm{Gamma}}(S_2|S_1, k) \quad (S_2 > 0) \end{cases} \qquad (S24)$$

Note that $\mathrm{P}(S_2)$ is discrete probability distribution for $S_2 = 0$ and continuous probability density for $S_2 > 0$. Using the same definition of $(\epsilon, \phi)$ for $s_1$ as binary models, $S_2$ distributions and corresponding channel capacity are fully determined in $(\epsilon, \phi)$ space. The evaluation of the $S_2$-channel capacity mapped in this space will allow the characterization of continuous models. As shown in **Fig. 10B, C**, there is no qualitative difference: both high $\epsilon$ and low $\phi$ are required for certain levels of performance, and large $\mathrm{n_{eff}}$ (about 5-6 for $C = 0.9$) is necessary at optimal $\epsilon$ of about 0.2. Only a quantitative difference was observed: the required $\phi$ gets larger approximately by a factor of $\alpha = 10$ compared with binary case, essentially because agonist binding events produce $\alpha$-times more $s_2$ than self ligands. Therefore, the advantage of multi-thread scheme is robustly seen.

**Appendix 8. Extended discussion on the cooperative firing behavior**

Cooperative firing is essential for the multi-thread scheme, and the fluctuation-driven nucleation model suggests that LAT condensation expectedly serves this step. Moreover, several other events in TCR signaling may also contribute to the non-linearity of firing step. For example, it has been suggested that Zap70 can be activated by intermolecular auto-phosphorylation (12), which suggests cooperative enhancement of Zap70 activity among completed threads. Grb2 (an adaptor protein) binding affinity to phosphorylated tyrosine residues of LAT is allosterically regulated, which also provides a source of cooperativity (13). Within each ITAM domain, two tyrosine residues get phosphorylated and a Zap70 molecule binds through avidity (14, 15). This corresponds to 2-thread-1-step scheme, where the avidity serves non-linearity, and there can be nested multi-thread structures within and among ITAM domains. More broadly, multiplicity-cooperativity combination, a ubiquitous structure in biochemical phenomena, has mostly been discussed in equilibrium conditions, and its distinct behaviors in non-equilibrium contexts are only recently getting recognized (16). The present study can be understood as novel non-equilibrium behaviors of this structure.

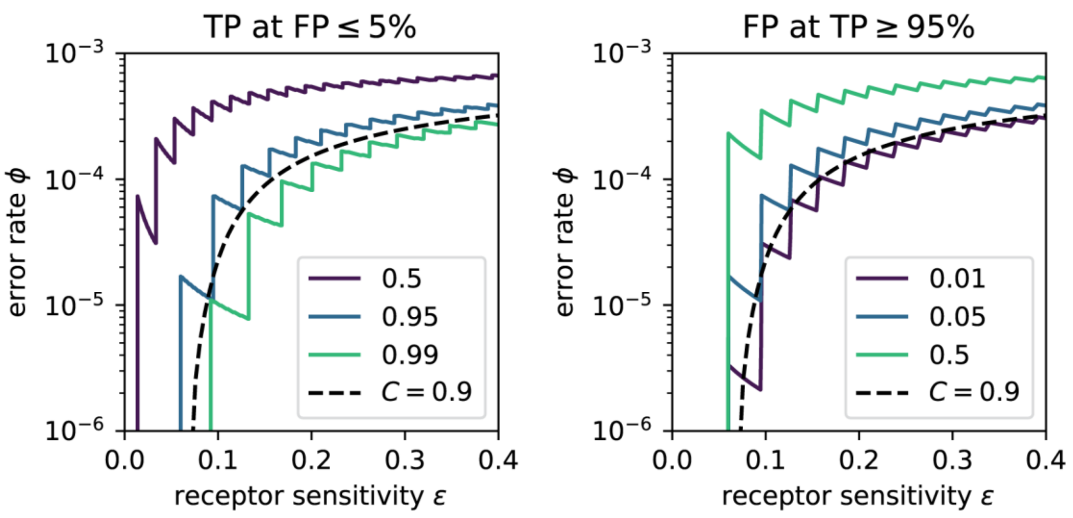


**Fig. S1.** TP-FP relationship as a function of receptor response. (left) The activation threshold $S_c$ is set to keep FP less than or equal to 5%, and the highest possible TP is shown as contours. (right) The activation threshold $S_c$ is set to keep TP greater than or equal to 95%, and the lowest possible FP is shown as contours. Note that the contours are discontinuous due to discrete $S$ distributions. The curve of $C = 0.9$ is overlaid.

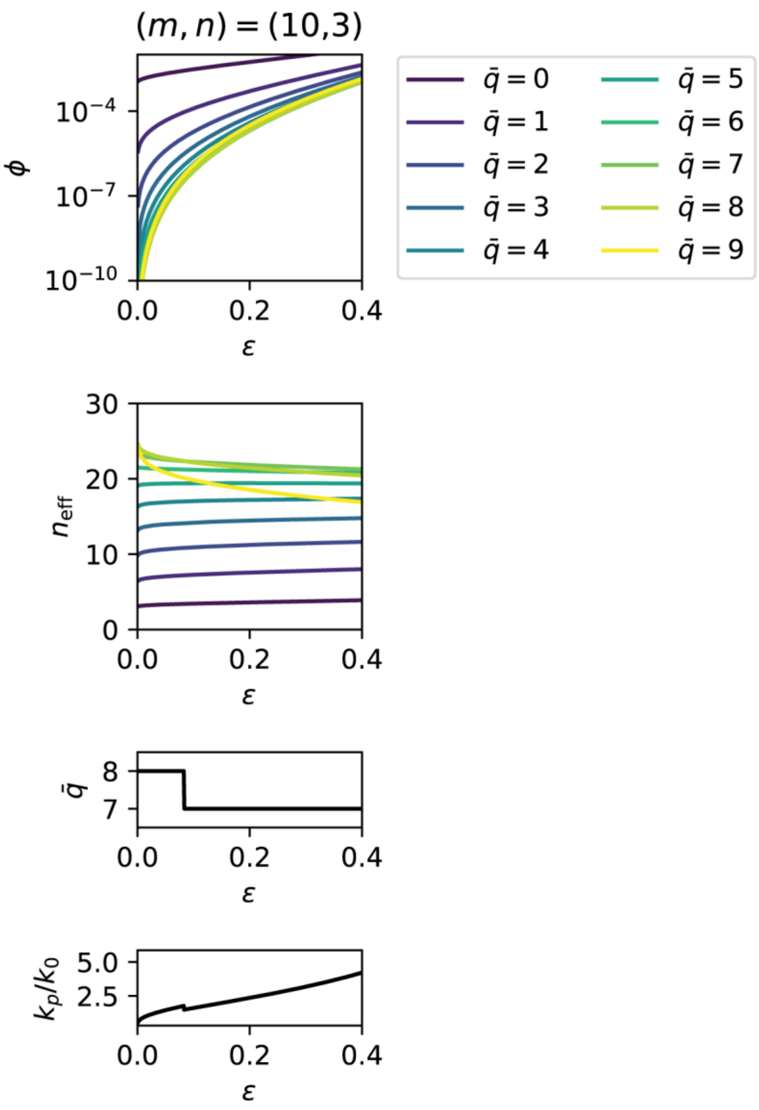


**Fig. S2.** An example of the parameter optimization for multi-thread scheme in a sharp firing regime. (top two) Receptor response and effective number of steps were calculated for all $\bar{q}$, and the trajectories are drawn. The lower envelope for $\phi$ and upper envelope for $n_{\mathrm{eff}}$ represents the trajectory with optimal parameters. (bottom two) The optimal parameter pair as a function of $\epsilon$. $k_p$ is shown in the unit of $k_0$.

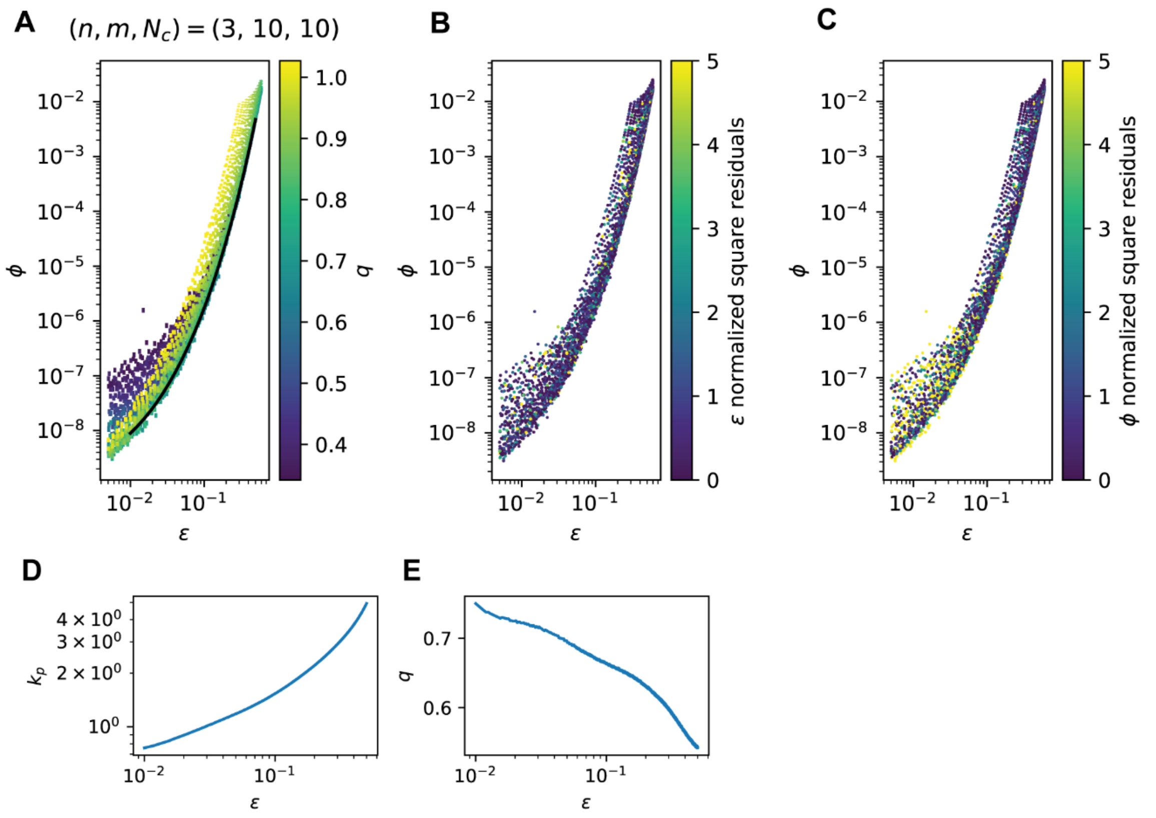


**Fig. S3.** An example of parameter optimization for multi-thread scheme with finite cooperativity exponent. (**A**) $(\epsilon, \phi)$ is calculated for an array of parameters $(k_p, q)$. Each datapoint has error bars in both axes representing the uncertainties from Monte-Carlo calculations. The lower envelope (black curve) is numerically obtained by fitting the empirical 2D spline (see Methods). (**B, C**) Fitting qualities were confirmed by inspecting the normalized square residuals. (**D, E**) Optimal paired parameters $(k_p, q)$ corresponding to the lower envelope are numerically determined.

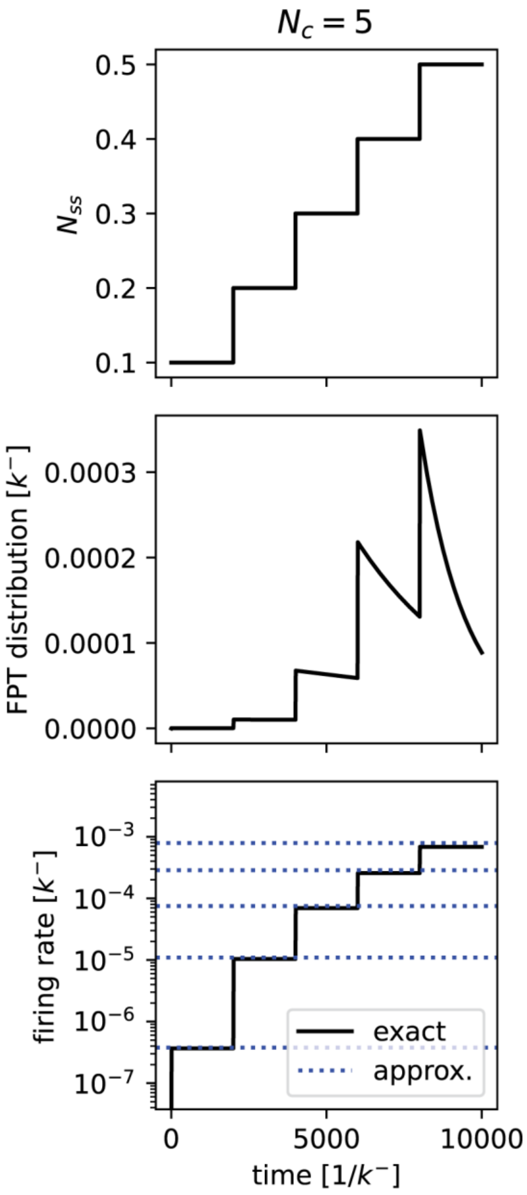


**Fig. S4.** Numerical calculation of LAT condensate nucleation kinetics. (top) $N_{ss}$ was varied over time in stepwise, reflecting stepwise increase of Zap70 level $r$. (middle) Calculated first passage time distribution by numerically solving the master equations. (bottom) Numerically calculated firing rate in the unit of $k^-$ ("exact"). Analytical solution from pseudo-equilibrium approximation for each $N_{ss}$ value is overlaid ("approx.").

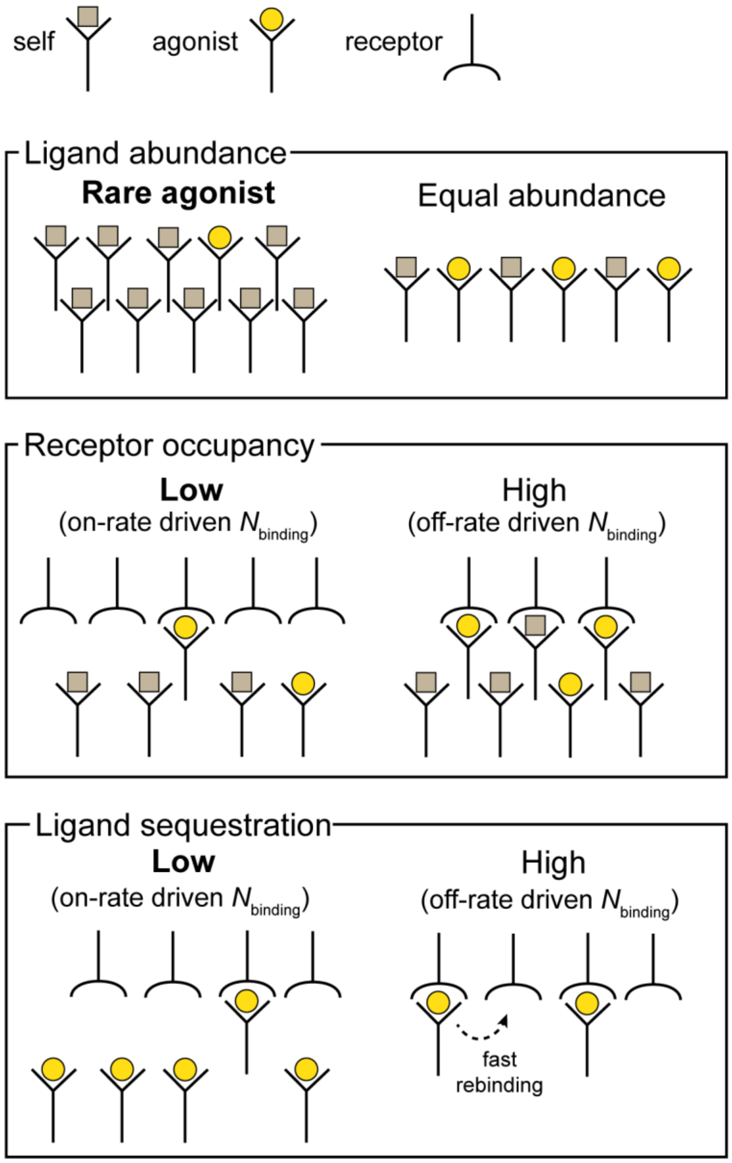


**Fig. S5.** Schematics of the assumptions influencing the number of binding events.

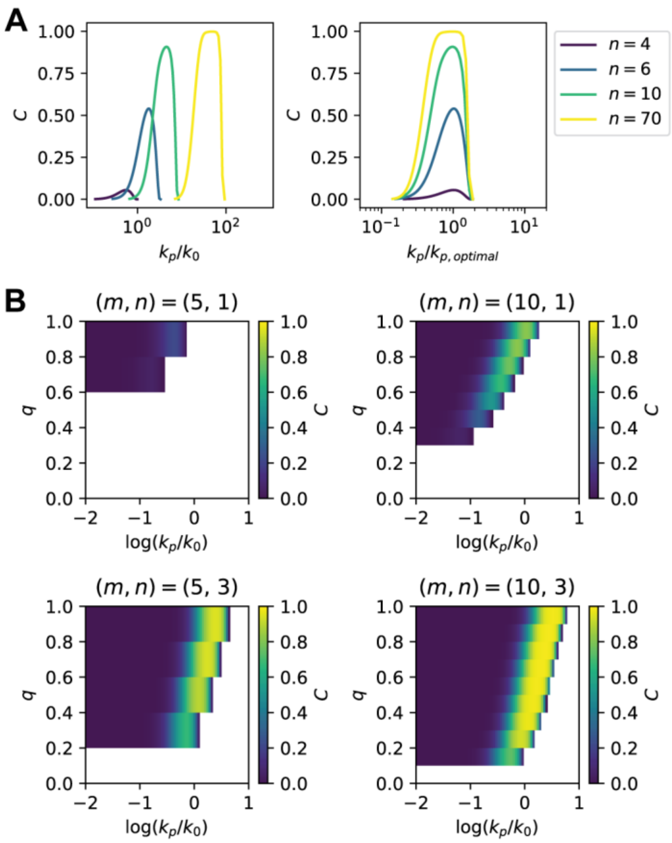


**Fig. S6.** Parameter-performance relationships for sequential scheme and multi-thread scheme in a sharp firing regime. (**A**, left panel) Discrimination performance $C$ as a function of $k_p$ (in the unit of $k_0$) for sequential scheme with different numbers of steps $n$. (right panel) $k_p$ is rescaled with respect to the optimal value of $k_p$ for each $n$. (**B**) Discrimination performance $C$ as a function of $k_p$ (in the unit of $k_0$, common logarithm) and $q$ for multi-thread scheme in a sharp firing regime with different $(m, n)$. The regime with $\beta\phi < 1$, for which self-induced signals exceed agonist-induced signals, is shown as blank.

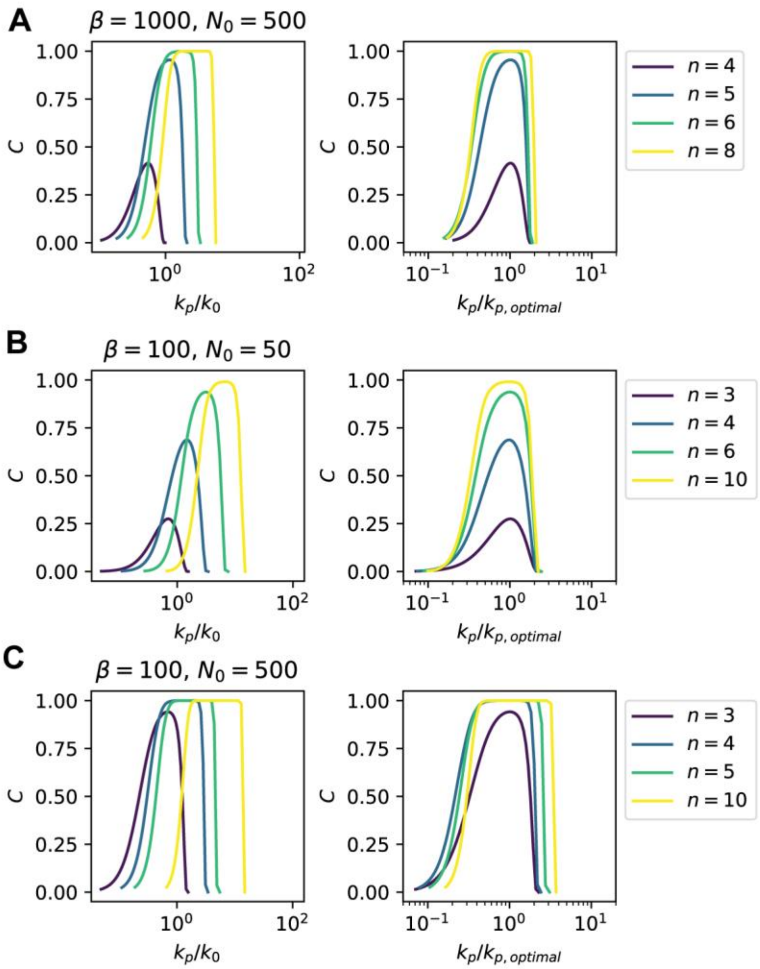


**Fig. S7.** Parameter-performance relationships for sequential scheme under different conditions for the number of binding events β and $N_0$.

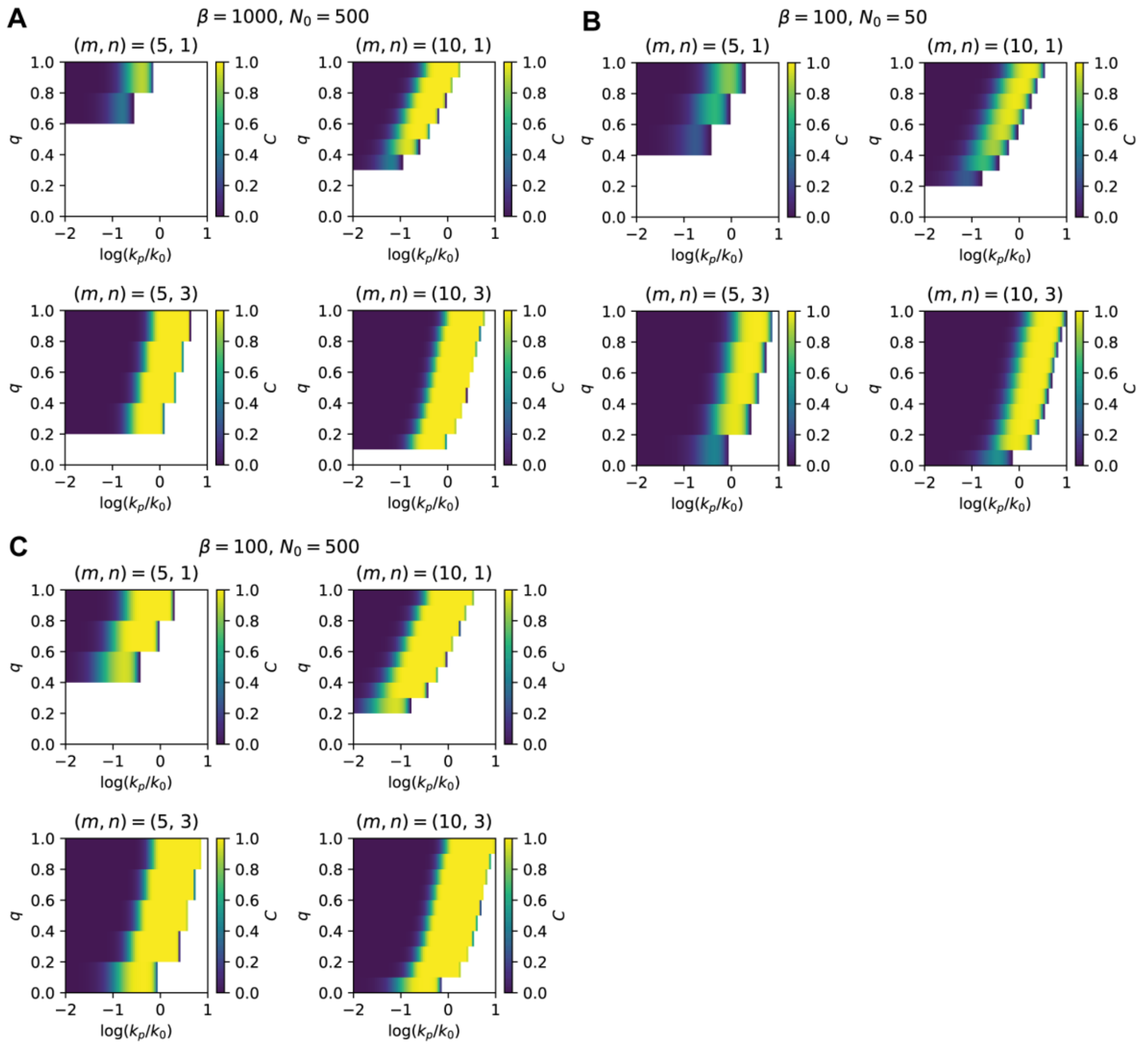


**Fig. S8.** Parameter-performance relationships for multi-thread scheme in a sharp firing regime under different conditions for the number of binding events β and $N_0$. The regime with $\beta\phi < 1$, for which self-induced signals exceed agonist-induced signals, is shown as blank.

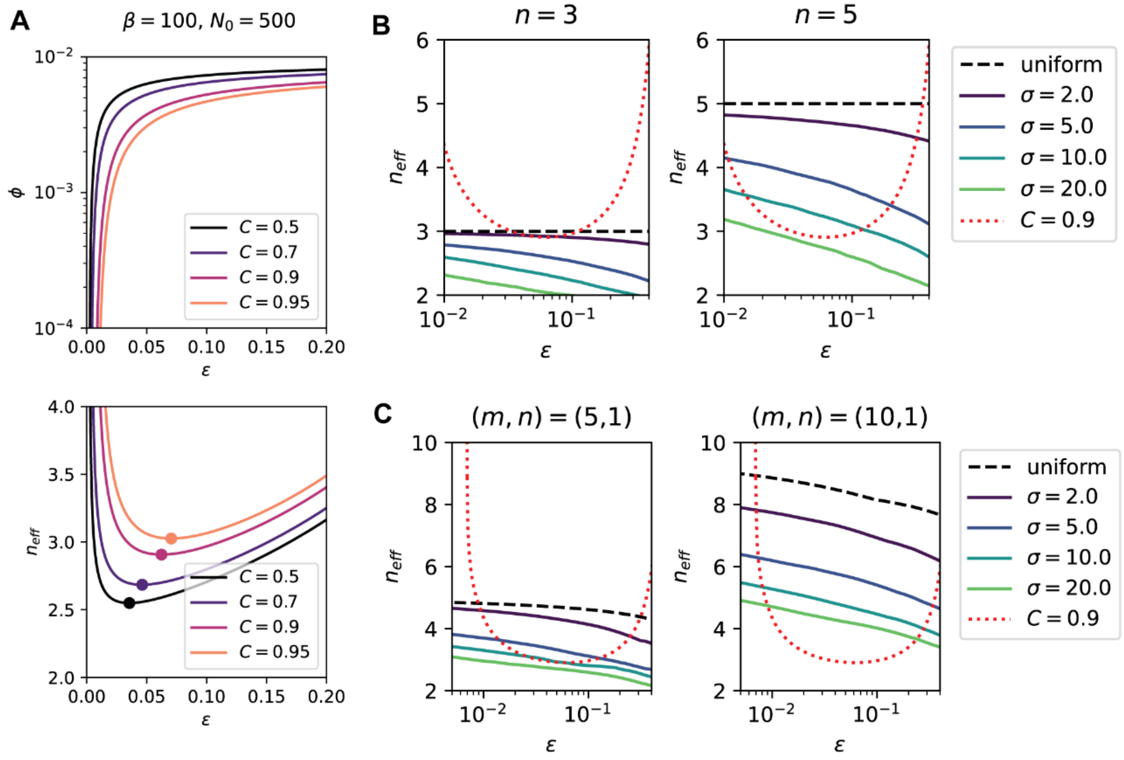


**Fig. S9.** Discrimination performance for low-noise condition. (**A**, top) Performance $C$ mapped in $(\epsilon, \phi)$ space under the condition $\beta = 100$, $N_0 = 500$. (bottom) The effective number of steps for corresponding $C$ values. Minimums are highlighted by markers. (**B**) Effective number of steps for sequential scheme with heterogeneous forward reaction rates. $\sigma$ is relative variation as described in Methods. The curve for $C = 0.9$ under the condition $\beta = 100$, $N_0 = 500$ is overlaid. (**C**) Effective number of steps for multi-thread scheme in a sharp firing regime and heterogeneous forward reaction rates in m-axis.

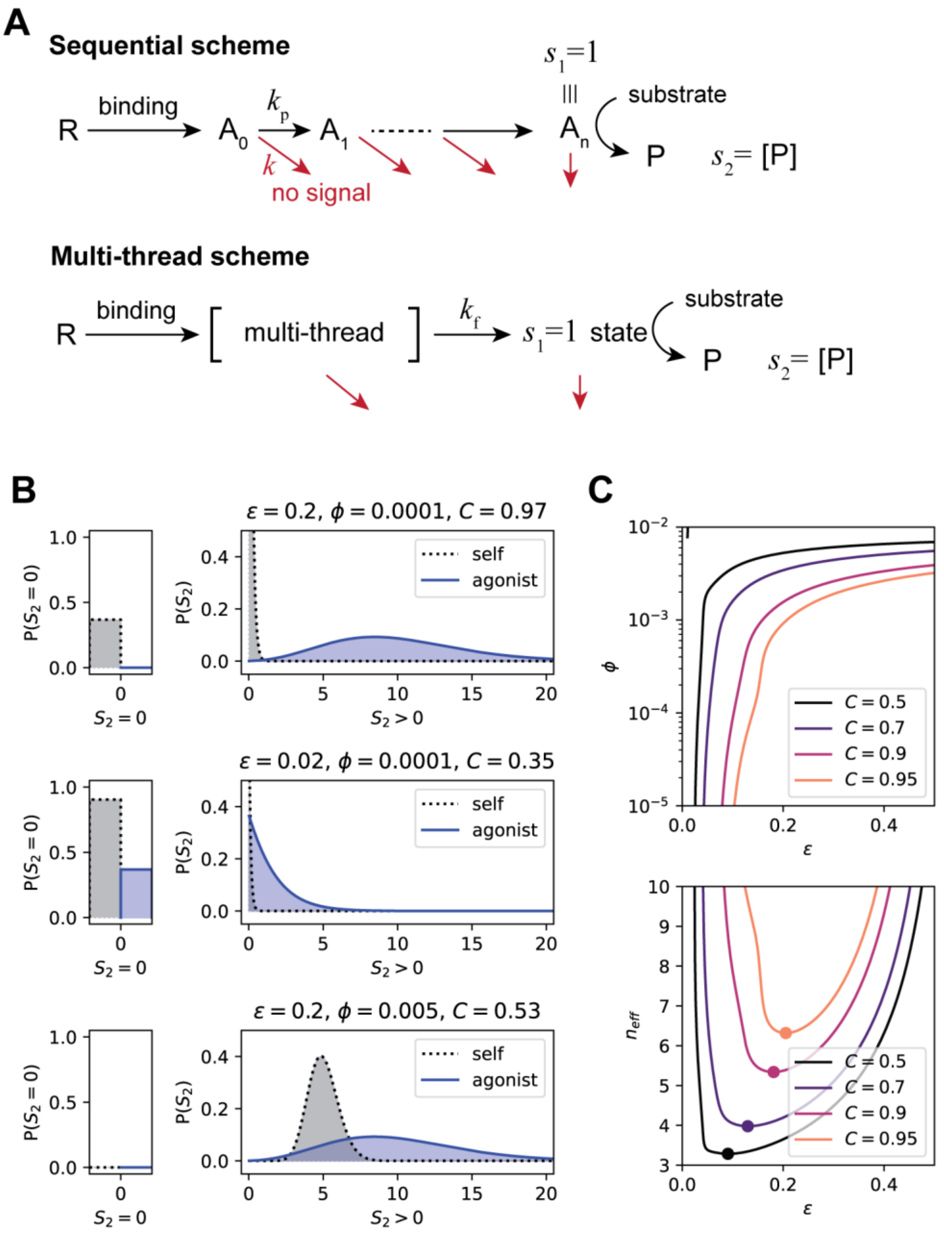


**Fig. S10.** Continuous-signal models. (**A**) Schematics for the continuous-signal models for sequential scheme and multi-thread scheme, respectively. (**B**) Distributions of sum continuous signal $S_2$ for representative $(\epsilon, \phi)$ pairs. Top: good performance with high receptor sensitivity and low error rate. Middle: poor performance with low receptor sensitivity. Bottom: poor performance with high error rate. (**C**) Discrimination performance $C$ defined for sum continuous signal $S_2$, and corresponding effective number of steps.
